# IL-17A-driven Macrophage Disorder Promotes Plaque Instability in Psoriatic Atherosclerosis

**DOI:** 10.1101/2025.05.21.654491

**Authors:** Canbing Dong, Jie Wang, Jui-Ming Lin, Xiaonian Lu, Junhao Zhu, Lanmei Lin, Lin Zhou, Fang Liu, Dan Wu, Xinyi Zhu, Jiazheng Sun, Jinhua Xu, Ningwen Zhu, Wenyu Wu, Jinyun Tan, Juan Du

## Abstract

Psoriasis is a common chronic inflammatory skin disease that predisposes individuals to cardiovascular diseases such as atherosclerosis. However, the underlying mechanisms linking these conditions have not yet been fully elucidated. Recent studies have indicated that the inflammatory cytokine IL-17A plays a significant role in both diseases. The aim of this study was to investigate how psoriasis affects the stability of atherosclerotic plaques via the IL-17A/IL-17RA pathway. We performed single-cell sequencing on patients with both psoriasis and atherosclerosis to analyze changes in the immune microenvironment within their carotid plaques and psoriatic lesions. We found that IL-17A significantly upregulated the expression of pro-inflammatory factors (e.g., CXCL8) in macrophages of psoriatic atherosclerosis patients, leading to weakening of the fibrous cap structure of plaques. Additionally, IL-17A further exacerbated plaque instability by interfering with macrophage lipid metabolism, particularly sphingolipid metabolism. Finally, using in vitro experiments and animal models, we demonstrated that IL-17A enhances pro-inflammatory and metabolic abnormalities in macrophages by downregulating PPARγ. Therapeutic strategies targeting IL-17A may contribute to the simultaneous mitigation of psoriasis and atherosclerosis-associated cardiovascular risk, providing new directions for clinical intervention.

## INTRODUCTION

Psoriasis is an immune-mediated systemic inflammatory skin disease often associated with various cardiovascular diseases ^1^. It is now considered an independent risk factor for cardiovascular disease, especially in patients with moderate-to-severe psoriasis, who exhibit a significantly increased risk (up to 40%) ^2^. Atherosclerosis is an inflammatory disease involving large and small arteries, and plaque rupture and thrombosis due to unstable plaques are the main causes of myocardial infarction and stroke ^3^. However, certain treatment regimens for severe psoriasis, such as acitretin and cyclosporine, can lead to hyperlipidemia, which adversely affects atherosclerosis^4,5^.

The specific mechanisms by which psoriasis leads to atherosclerotic plaque destabilization remain to be thoroughly investigated.

During the psoriasis-induced aggravation of atherosclerosis, remodeling of the plaque immune microenvironment plays a key role in plaque destabilization.

Macrophages, considered the core cells of the plaque immune microenvironment ^6, 7^, are further activated in plaques under the stimulation of psoriatic inflammatory factors such as TNF-α, IL-17, IL-23, and IFN-γ. Activated macrophages secrete inflammatory cytokines (TNF-α, IL-1β, and IL-6) and chemokines (CXCL1, CXCL2, and CXCL8), which recruit neutrophils and T cells, leading to amplified inflammation and plaque instability ^8–11^. Furthermore, macrophages produce large amounts of matrix metalloproteinases (e.g., MMP-2/9), which degrade collagen fibers in the plaque fibrous cap, leading to thinning of the fibrous cap and increasing the risk of plaque rupture ^12–14^. We recently reported upregulation of MMP2, an important plaque-destabilizing factor ^15–18^, in the peripheral blood of patients with moderate-to-severe plaque psoriasis ^15^, suggesting a possible increased risk of plaque rupture due to psoriasis.

Among the inflammatory factors significantly increased in psoriasis, IL-17A is one of the most important cytokines in the pathogenesis and progression of the disease. Many biologics targeting IL-17A have been widely used in the clinic with significant results ^1^. Current studies suggest that IL-17A and its signaling pathway play a major role in atherosclerosis progression and plaque instability ^19–22^. IL-17A promotes the recruitment of inflammatory cells, such as macrophages and neutrophils, and maintains chronic inflammation in the arterial wall by regulating various inflammatory factors, including TNF-α, IL-6, IL-1β, and MCP-1 ^23, 24^. Additionally, IL-17A can induce macrophages to produce matrix metalloproteinases (MMPs) ^25, 26^. These proteases degrade collagen fibers in plaques, weakening the stability of the fibrous cap and making plaques more prone to rupture. However, IL-17A-mediated changes in the immune microenvironment in the context of psoriasis have not been elucidated.

To understand the mechanisms by which psoriasis alters the plaque immune microenvironment to mediate atherosclerotic plaque destabilization, we recruited patients with both carotid plaques and moderate-to-severe plaque-type psoriasis at Huashan Hospital. We collected their carotid plaques and psoriatic lesions for single-cell transcriptome sequencing. Based on complementary single-cell mapping of psoriasis-atherosclerosis tissues, we explored the specific mechanisms by which psoriasis, particularly IL-17A, mediates plaque progression and instability. We validated these mechanisms and phenomena through in vivo and ex vivo experiments.

## RESULTS

### Establishment of Single-Cell Mapping of Plaques and Lesions in Psoriatic Atherosclerosis Patients

To investigate the specific mechanisms by which psoriasis and atherosclerosis interact, we recruited patients with both conditions, patients with psoriasis only, and patients with atherosclerosis only. From psoriatic atherosclerosis patients who met the indications for carotid plaque surgery, we collected psoriatic lesions and carotid plaques for single-cell transcriptome sequencing. Control samples included lesions from psoriasis-only patients and plaques from atherosclerosis-only patients (Fig. 1a).

**Fig. 1.**
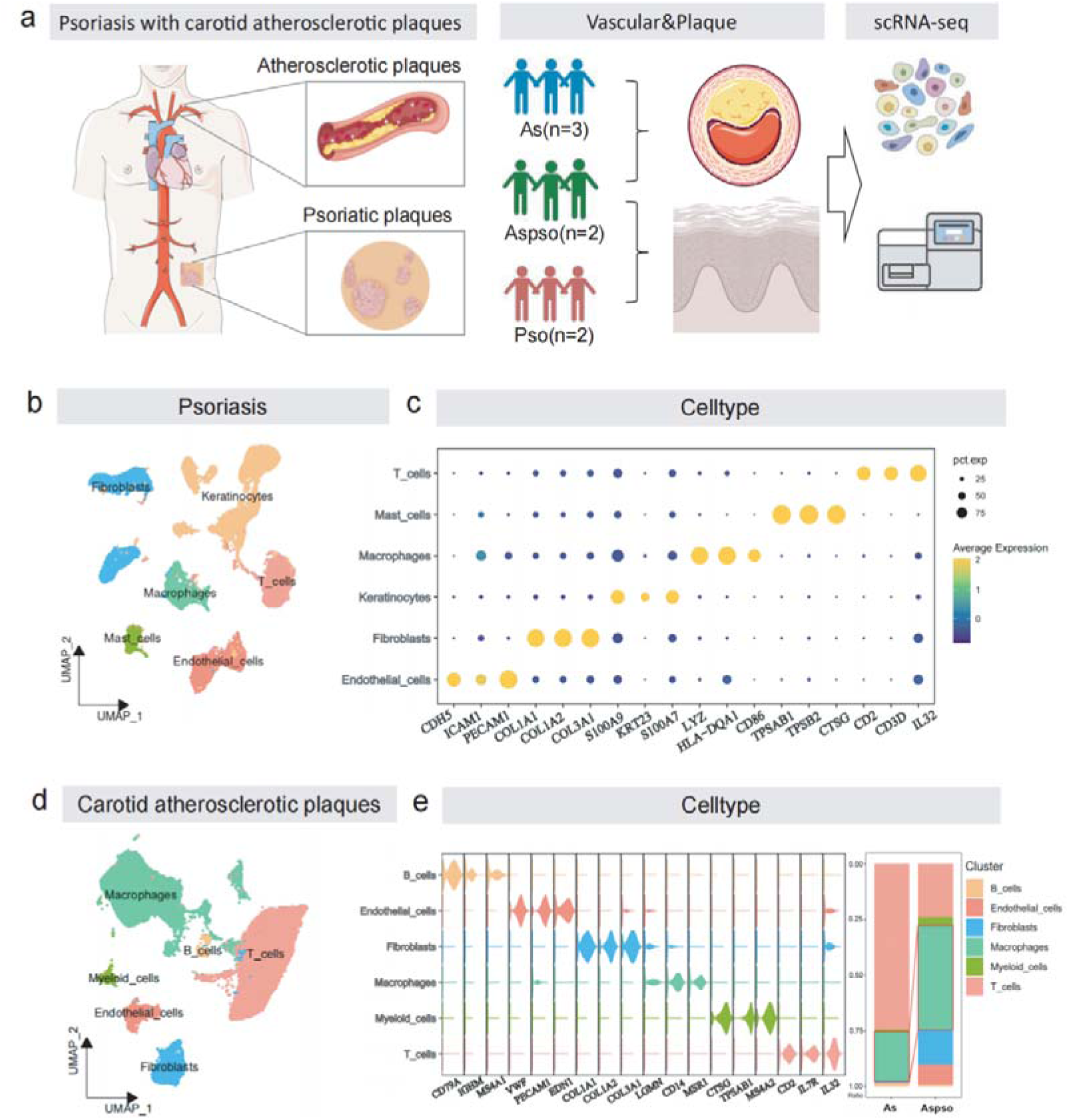
Single-Cell Mapping of Psoriasis-Atherosclerosis Co-Morbidities. **a,** Schematic representation of the study design. Psoriatic lesions and carotid plaques were collected from patients with both moderate-to-severe plaque psoriasis and carotid atherosclerosis and subjected to single-cell transcriptome sequencing. **b,** Uniform Manifold Approximation and Projection (UMAP) demonstrating the cellular classification of psoriatic lesions in co-morbid patients and controls. **c,** Basis for cell classification in psoriatic lesions. **d,** UMAP showing cellular classification of plaques in co-morbid patients and controls. **e,** Basis for cell classification in atherosclerotic plaques and the percentage of cells in each category. Aspso: Patients with both psoriasis and carotid plaques; Pso: Patients with psoriasis only; As: Patients with carotid plaques only.

Uniform Manifold Approximation and Projection (UMAP) and cell type annotation in psoriatic skin lesions identified six cell types: endothelial cells, keratinocytes, fibroblasts, mast cells, macrophages, and T cells (Fig. 1b). Macrophages were identified using markers LYZ, HLA-DQA1, and CD86; fibroblasts with COL1A1, COL1A2, and COL3A1; and endothelial cells with CDH5, ICAM1, and PECAM1 (Fig. 1c). Similarly, all cells in the plaques were categorized into myeloid cells, endothelial cells, macrophages, T cells, fibroblasts, and B cells (Fig. 1d).

Macrophage markers included LGMN, CD14, and MSR1; fibroblast markers were COL1A1, COL1A2, and COL3A1; and endothelial cell markers were VWF, PECAM1, and EDN1 (Fig. 1e). Analysis of cellular composition within plaques revealed a significantly higher percentage of macrophages in atherosclerotic plaques from psoriasis patients compared to those from non-psoriatic atherosclerosis patients (Fig. 1e), suggesting that psoriasis may influence the progression of atherosclerosis through macrophages.

### Pro-Inflammatory Interactions of Macrophages with Endothelial Cells and Fibroblasts in the Immune Microenvironment of Co-Morbid Plaques and Lesions

We used CellChat to probe the interaction signals between different cells in the plaques to explore macrophage interactions with various cells in psoriatic atherosclerosis plaques. The results, as shown in Fig. 2a, suggest that macrophages have a greater number of cellular interactions and higher interaction weights with fibroblasts and endothelial cells. Further analysis of specific interaction signals showed that the CXCL2/3/8–ACKR1 interaction pathway was heavily upregulated in macrophages interacting with endothelial cells (Fig. 2b). Among the various chemokine signals in macrophages, CXCL8 was most significantly upregulated (Fig. 2c). Macrophage chemokine expression was verified by single-cell sequencing in psoriatic atherosclerosis plaques (Aspso) and atherosclerosis-only plaques (As), as shown in Fig. 2d; CXCL8, CXCL2, CXCL3, CCL2, and CCL3 were significantly upregulated in psoriatic atherosclerosis plaques.

**Fig. 2.**
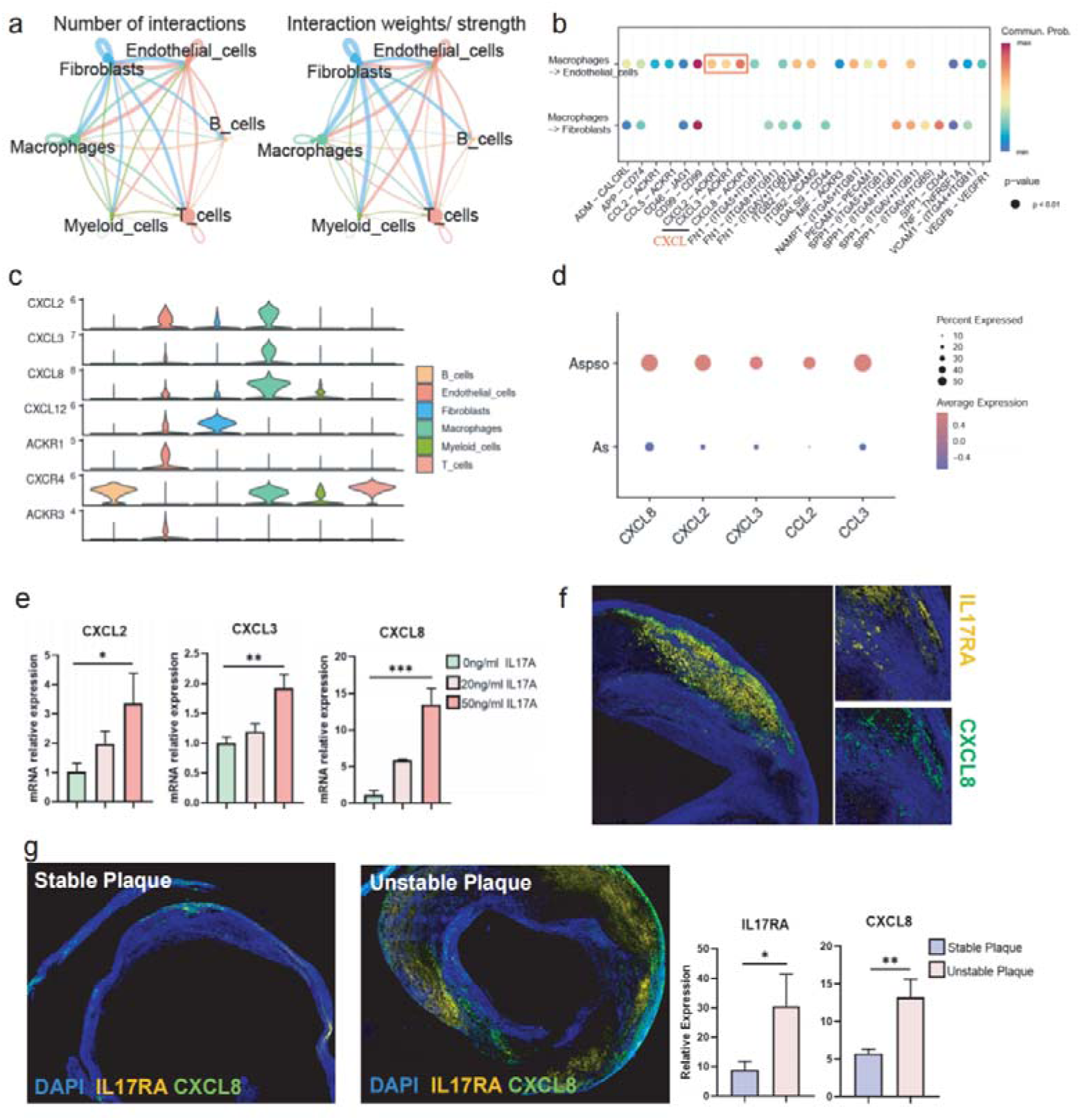
Macrophages in Co-Morbid Plaques Produce More Chemokines. **a,** Number of interactions and interaction weights among each cell type in plaques. **b,** Interaction signals between macrophages and endothelial cells and fibroblasts in plaques. **c,** Expression levels of chemokine signals in various cell types within plaques. **d,** Comparison of chemokine expression levels in co-morbid and control plaques. **e,** mRNA expression levels of CXCL2, CXCL3, and CXCL8 in macrophages after IL-17A intervention. **f,** Immunofluorescence showing co-localization of IL-17RA and CXCL8 in co-morbid plaques. **g,** Immunofluorescence showing the expression of IL-17RA and CXCL8 in stable and unstable plaques. Significance indicated as *P < 0.05, **P < 0.01, ***P < 0.001, with ns denoting not significant. Statistical analysis was performed using the One-way ANOVA.

IL-17A is an important pro-inflammatory factor, and we treated macrophages with IL-17A recombinant proteins; qPCR revealed that the corresponding chemokines CXCL2, CXCL3, and CXCL8 were significantly upregulated (Fig. 2e).

Immunofluorescence co-localization revealed that IL-17RA is accompanied by upregulation of CXCL8 (Fig. 2f). CXCL8 is the most potent neutrophil chemokine known to direct neutrophils toward plaque sites. The aggregation of neutrophils increases the local inflammatory response, while the enzymes they release disrupt the matrix, thinning the plaque fibrous cap and promoting plaque instability ^8, 9^. We collected unstable and stable plaques for immunofluorescence, and the results suggested that IL-17RA and CXCL8 were more widely distributed in unstable plaques (Fig. 2g). Information on the collected stable and unstable plaques is shown in Table S1 and was confirmed by Masson staining (Extended Data Fig1a). This suggests that IL-17A may promote the instability of atherosclerotic plaques through CXCL8.

The inflammatory properties of endothelial cells and fibroblasts likewise play important roles in the development and progression of atherosclerosis. The substantial upregulation of CXCL8–ACKR1 interactions between macrophages and endothelial cells has been found previously, and our secondary fractionation of endothelial cells in plaques revealed that ACKR1 was significantly upregulated in psoriatic atherosclerosis plaques (Extended Data Fig1b). Differential genes between ACKR1 high- and low-expressing endothelial cells were screened for enrichment analysis. GO enrichment analysis revealed involvement in the positive regulation of inflammatory response (Extended Data Fig1c). Upon treatment of endothelial cells with recombinant IL-17A protein, qPCR confirmed the upregulation of ACKR1 and the endothelial inflammation marker ICAM1 mRNA (Extended Data Fig1d). Similarly, analysis of differential genes in fibroblasts in psoriatic atherosclerosis plaques and controls showed upregulation of inflammatory factors such as MMP2, MMP14 and VCAN in psoriatic atherosclerosis plaques (Extended Data Fig1e). These findings suggest upregulation of endothelial cell ACKR1 and heightened pro-inflammatory properties of fibroblasts in psoriatic atherosclerosis plaques.

Given the close association between CXCL8 and neutrophils, we performed a secondary analysis focusing on myeloid cells within the plaques (Extended Data Fig2a). The results showed that, compared to conventional plaques (As), comorbid plaques (Aspso) exhibited significantly higher expression of CXCL8 receptors CXCR2 and ACKR1 (Extended Data Fig2b). Differential gene analysis between the Aspso and As groups revealed that GO enrichment was significantly associated with myeloid cell stabilization, differentiation, and activation (Extended Data Fig2c).

Among the genes involved in these enriched pathways, MMP9 was markedly upregulated in the Aspso group (Extended Data Fig2d).

Psoriatic lesions of psoriatic atherosclerosis patients similarly demonstrated an enhanced pro-inflammatory phenotype of macrophages, endothelial cells, and fibroblasts. Using CellChat to explore interaction signals between different cells in skin lesions, we likewise found that macrophages and endothelial cells have abundant cellular interactions in both number and weight (Extended Data Fig3a). Among the specific interaction signals, CXCL2/8–ACKR1 interactions between macrophages and endothelial cells were still present (Extended Data Fig3b), with CXCL8 being significantly upregulated (Extended Data Fig3c). Immunofluorescence showed increased expression of macrophages and CXCL8 in co-morbid lesions, supporting these findings (Extended Data Fig3d). Our time-series analysis of psoriatic skin lesions from psoriatic atherosclerosis patients revealed that macrophages showed concomitant upregulation of CXCL8 expression with the upregulation of the IL-17A receptor IL-17RA (Extended Data Fig3e), and immunofluorescence verified the co-localization of CXCL8 and IL-17RA (Extended Data Fig3f). These results suggest that IL-17A likewise promotes upregulation of macrophage CXCL8 expression and inflammation in psoriatic lesions.

Endothelial cells and fibroblasts were also analyzed in depth. Endothelial cells were divided into two groups based on high and low ACKR1 expression, and differential genes were screened for enrichment analysis (Extended Data Fig4a,b). GO enrichment analysis revealed involvement in epidermal proliferation and migration, neutrophil migration, and other pathways associated with psoriasis pathogenesis (Extended Data Fig4c). Similarly, screening for differential genes in fibroblasts from co-morbid skin lesions and controls showed upregulation of inflammatory markers MMP2, CCL2, and CXCL12 in co-morbid skin lesion fibroblasts (Extended Data Fig4d). These results suggest a promotional effect of IL-17A on the upregulation of ACKR1 in skin lesion endothelial cells and increased inflammatory properties of fibroblasts. In conclusion, IL-17A promotes the production of macrophage chemokines, especially CXCL8, which mediates the progression of both atherosclerosis and psoriasis.

### IL-17A Promotes Atherosclerotic Plaque Instability by Regulating Macrophage Chemokine and Lipid Metabolism

To investigate the specific effects of IL-17A on macrophages in plaques, we divided macrophages into IL17RA_hi and IL-17RA_lo groups and performed differential gene expression analysis (Extended Data Fig5a). Gene Ontology (GO) enrichment analysis showed that the differentially expressed genes were enriched in pathways related to chemokine production, neutrophil recruitment, proliferation, adhesion, and migration (Fig. 3a), consistent with our previous finding of upregulated neutrophil-recruiting chemokines such as CXCL2 and CXCL8 in psoriatic atherosclerosis plaques. Kyoto Encyclopedia of Genes and Genomes (KEGG) enrichment analysis revealed close associations with the IL-17 signaling pathway, lipid and atherosclerosis, chemokine signaling, sphingolipid signaling pathway, and PPAR signaling pathway (Fig. 3b). This suggests that IL-17A is involved in macrophage pro-inflammatory differentiation and intracellular lipid metabolism processes.

**Fig. 3.**
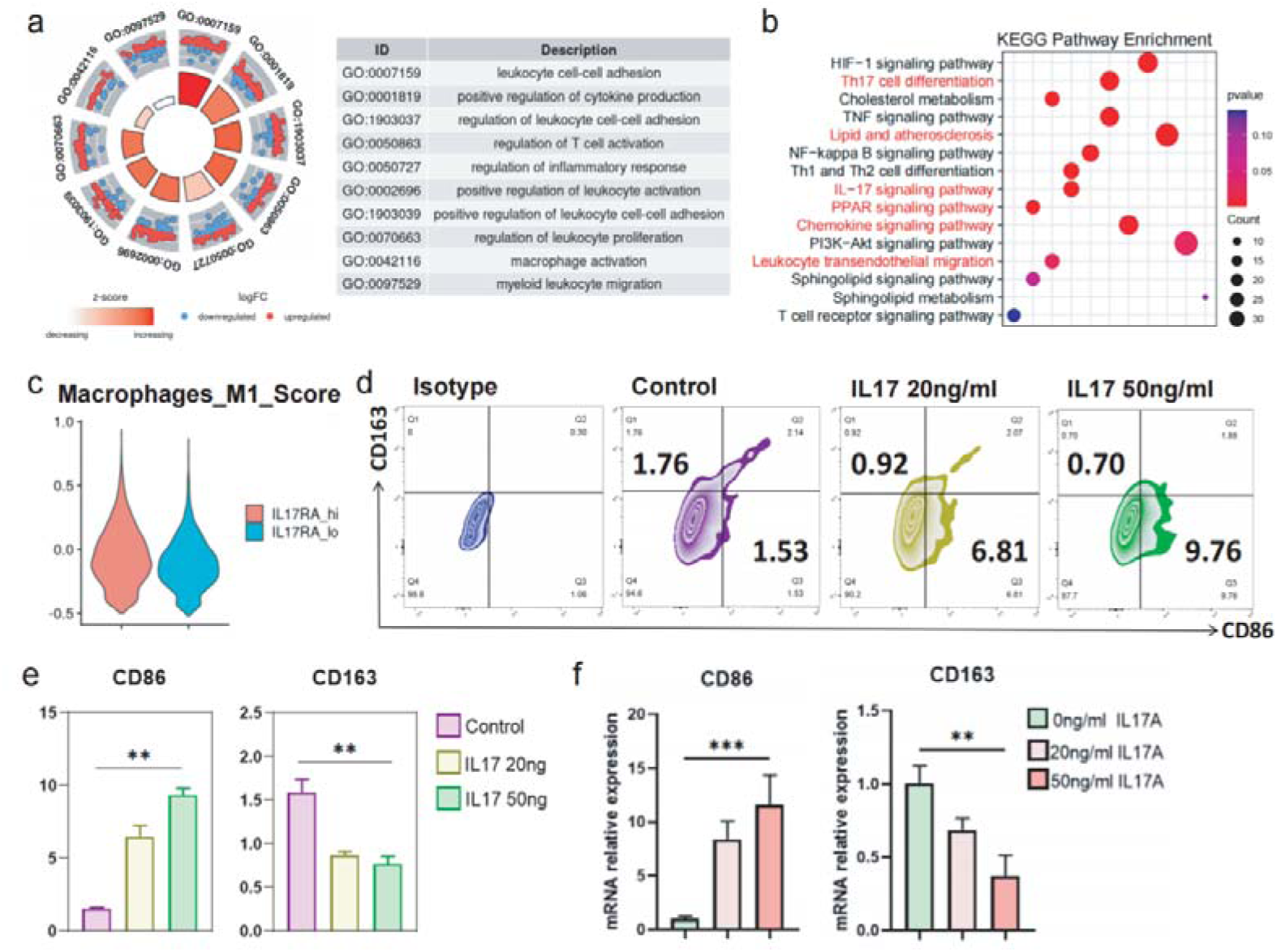
IL-17A Upregulates Macrophage Inflammation. **a,** Gene Ontology (GO) enrichment analysis of differential genes in macrophages from co-morbid versus control plaques. **b,** Kyoto Encyclopedia of Genes and Genomes (KEGG) enrichment analysis of differential genes in macrophages from co-morbid versus control plaques. **c,** M1 polarization gene set scoring in macrophages from IL17RA_hi and IL17RA_lo plaques. **d,** Flow cytometry showing upregulation of CD86 and downregulation of CD163 in macrophages after IL-17A intervention. **e,** Statistical analysis of three independent experimental replicates. **f,** mRNA expression levels of CD86 and CD163 in macrophages after IL-17A intervention. Significance indicated as *P < 0.05, **P < 0.01, ***P < 0.001, with ns denoting not significant. Statistical analysis was performed using the One-way ANOVA and the t-test.

Classically, macrophages are divided into pro-inflammatory M1 type and anti-inflammatory M2 type. M1 macrophages secrete pro-inflammatory cytokines and matrix metalloproteinases (MMP2/9), promoting atherosclerosis progression and plaque destabilization ^12, 13^. Using AddModuleScore to assess M1 polarization of macrophages in the IL17RA_hi and IL-17RA_lo groups, we found enhanced M1 polarization in the IL17RA_hi group (Fig. 3c). We treated macrophages with 20 ng/mL and 50 ng/mL IL-17A recombinant protein, and flow cytometry showed that as the concentration of IL-17A increased, the M1 polarization marker CD86 was upregulated, while the M2 marker CD163 was downregulated (Fig. 3d,e). qRT-PCR similarly showed upregulation of the M1 marker (CD86) and downregulation of the M2 marker (CD163) (Fig. 3f). With advances in macrophage classification, recent studies have identified three subtypes: IL-1β□, LYVE1□, and TREM2□ macrophages, among which IL-1β□ macrophages are associated with plaque rupture. Our analysis revealed that macrophages in the IL-17RA_hi group expressed higher levels of IL-1β, whereas those in the IL-17RA_lo group expressed more TREM2 (Extended Data Fig5b).

IL-17A may mediate macrophage disruption by interfering with intracellular lipid metabolism. We treated macrophages with 50 ng/mL IL-17A recombinant protein and analyzed metabolite changes using liquid chromatography-tandem mass spectrometry (LC-MS/MS). As shown in Fig. 4a, IL-17A significantly downregulated lipid and lipid-like molecules involved in cholesterol metabolism, sphingolipid metabolism, phospholipid metabolism, and triglyceride metabolism. Among these, metabolites related to phospholipid metabolism (e.g., PE(18:0/18:2), DHA-PC, PG(16:0/18:1), PG(18:1/18:1(9Z)), PC(16:0/16:0), GPC, c9-PA, c11-EA, LPG(18:1), O-PEA) were most significantly downregulated. Enrichment analysis of differential metabolites also showed abnormalities in sphingolipid and glycerophospholipid metabolism (Fig. 4b,c). These results suggest that IL-17A may contribute to atherosclerotic plaque destabilization by downregulating macrophage sphingolipid and glycerophospholipid metabolites.

**Fig. 4.**
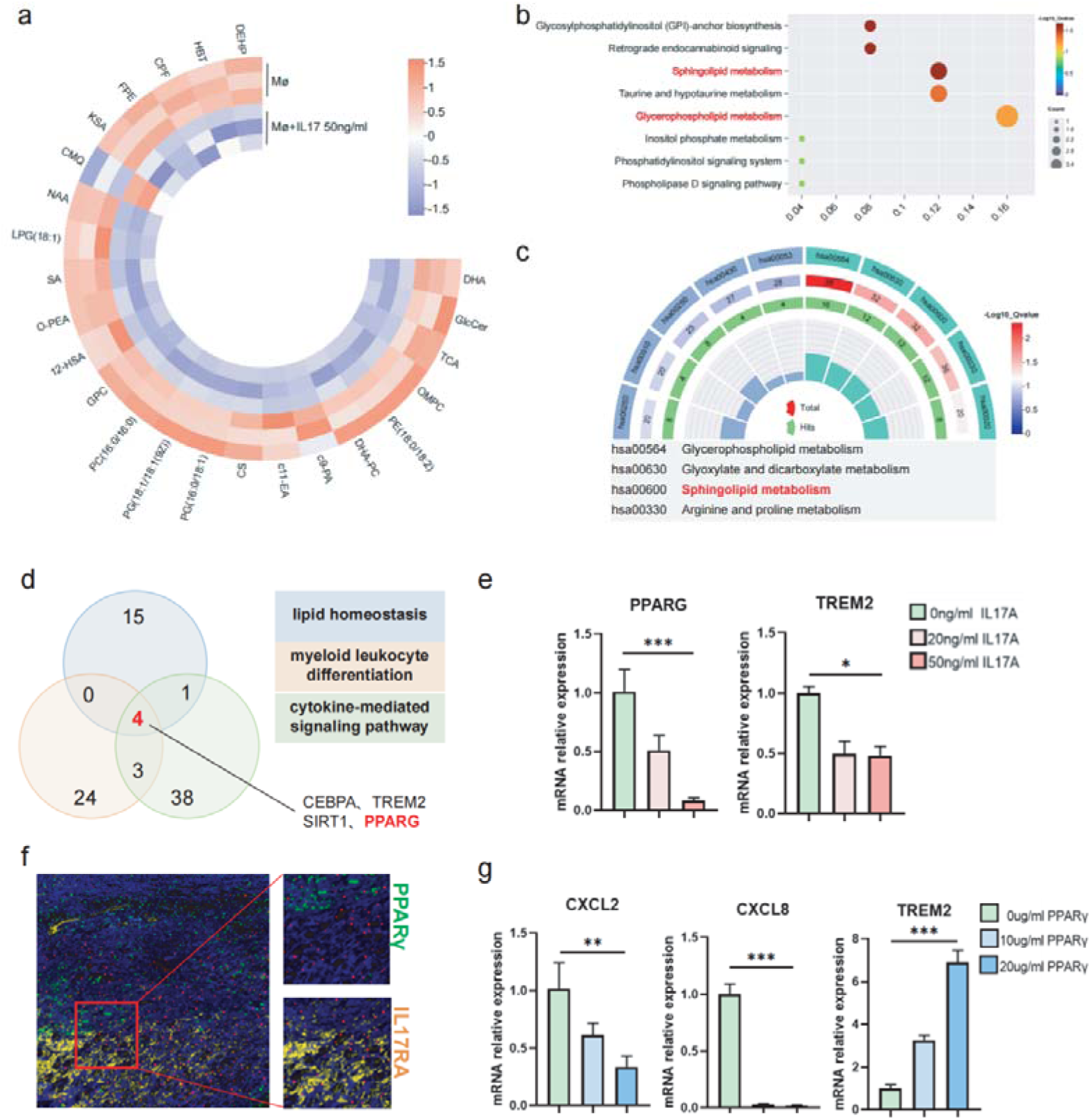
IL-17A Reduces Sphingomyelin Components in Macrophages by Downregulating PPAR_γ_. **a,** Changes in various lipid components in macrophages after IL-17A intervention. **b-c,** Enrichment analysis showing the main downregulated lipid types. **d,** Gene interaction network showing that PPARG is involved in three enrichment pathways: lipid homeostasis□cytokine-mediated signaling pathway□myeloid leukocyte differentiation. **e,** mRNA expression levels of PPARG, TREM2 in macrophages after IL-17A intervention. **f,** Immunofluorescence showing the spatial relationship between IL-17RA and PPARγ. **g,** mRNA expression levels of CXCL2, CXCL8, and TREM2 in macrophages after PPARγ agonist intervention. Significance indicated as *P < 0.05, **P < 0.01, ***P < 0.001, with ns denoting not significant. Statistical analysis was performed using the One-way ANOVA.

Similarly, we divided macrophages into Aspso and As groups and performed differential gene analysis followed by GO and KEGG enrichment analyses. The results similarly indicated enrichment in pathways related to T cell and neutrophil activation, IL-17 signaling, PPAR signaling, and sphingolipid metabolism (Extended Data Fig5c,d). AddModuleScore analysis also showed enhanced M1 polarization in the Aspso group (Extended Data Fig5e).

In conclusion, IL-17A may mediate the destabilization of atherosclerotic plaques by promoting macrophage production of inflammatory factors, recruiting neutrophils, and downregulating lipid metabolites. To further investigate the regulatory role of IL-17A on macrophages, we examined the intersection of three enrichment pathways: lipid homeostasis, cytokine-mediated signaling pathway, myeloid leukocyte differentiation (Fig. 4d). The results indicated that PPARG (PPARγ) were involved in all three pathways. KEGG enrichment analysis of differentially expressed genes between IL-17RA_hi and IL-17RA_lo macrophages also revealed involvement in the PPAR signaling pathway. Using IL-17A to treat macrophages, qRT-PCR showed significant downregulation of PPARG (PPARγ) (Fig. 4e). Spatial transcriptomics showed that sites of IL-17RA upregulation in plaques were accompanied by downregulation of PPARγ (Extended Data Fig5f), which was confirmed by immunofluorescence (Fig. 4f). Treating macrophages with 10 μg/mL and 20 μg/mL Pioglitazone (PPARγ agonists) resulted in downregulation of chemokines CXCL2, CXCL8, and upregulation of lipid metabolism genes TREM2 (Fig. 4g). These findings suggest that IL-17A may influence the inflammatory and lipid metabolic state of macrophages by downregulating PPARγ.

In psoriatic skin lesions, we similarly performed secondary clustering of macrophages, comparing and enriching for differential genes of IL17RA_hi and IL-17RA_lo groups (Extended Data Fig6a). GO enrichment analyses also showed recruitment of cytokine production, neutrophil chemotaxis, and other pathways (Extended Data Fig6b). KEGG enrichment analyses suggested enrichment in lipid metabolism and atherosclerosis, the IL-17 signaling pathway, chemokine signaling pathways, neutrophil migration, and PPAR signaling pathway (Extended Data Fig6c), consistent with plaque-related analyses. Overall, these results suggest that IL-17A may contribute to macrophage-mediated destabilization by downregulating PPARγ to promote macrophage inflammation, chemokine production, and reduction of intracellular sphingolipid components.

### PPAR**_γ_** Agonists Can Reverse Macrophage Disorganization Caused by IL-17A

To verify that PPARγ agonists can reverse the abnormalities of chemokines and metabolic status caused by IL-17A, we treated macrophages with 50 ng/mL IL-17A recombinant protein and, after one day, intervened with 10 μg/mL and 20 μg/mL Pioglitazone (PPARγ agonist). As shown by qRT-PCR and flow cytometry, the upregulation of the macrophage M1 polarization marker CD86 induced by IL-17A was reversed by PPARγ agonists (Fig. 5a,b). mRNAs of chemokines such as CCL2, CCL3, CXCL1, CXCL2, and CXCL8 were significantly upregulated after IL-17A treatment and showed gradient downregulation with PPARγ agonist intervention (Fig. 5c). Lipid metabolism-related mRNAs, such as TREM2, PLTP, APOE, FABP3 and FABP5, were downregulated after IL-17A treatment and upregulated after PPARγ agonist intervention (Fig. 5d). These results suggest that the M1 polarization, inflammatory cytokine secretion, and lipid metabolism disorders in macrophages caused by IL-17A can be reversed by PPARγ agonists.

**Fig. 5.**
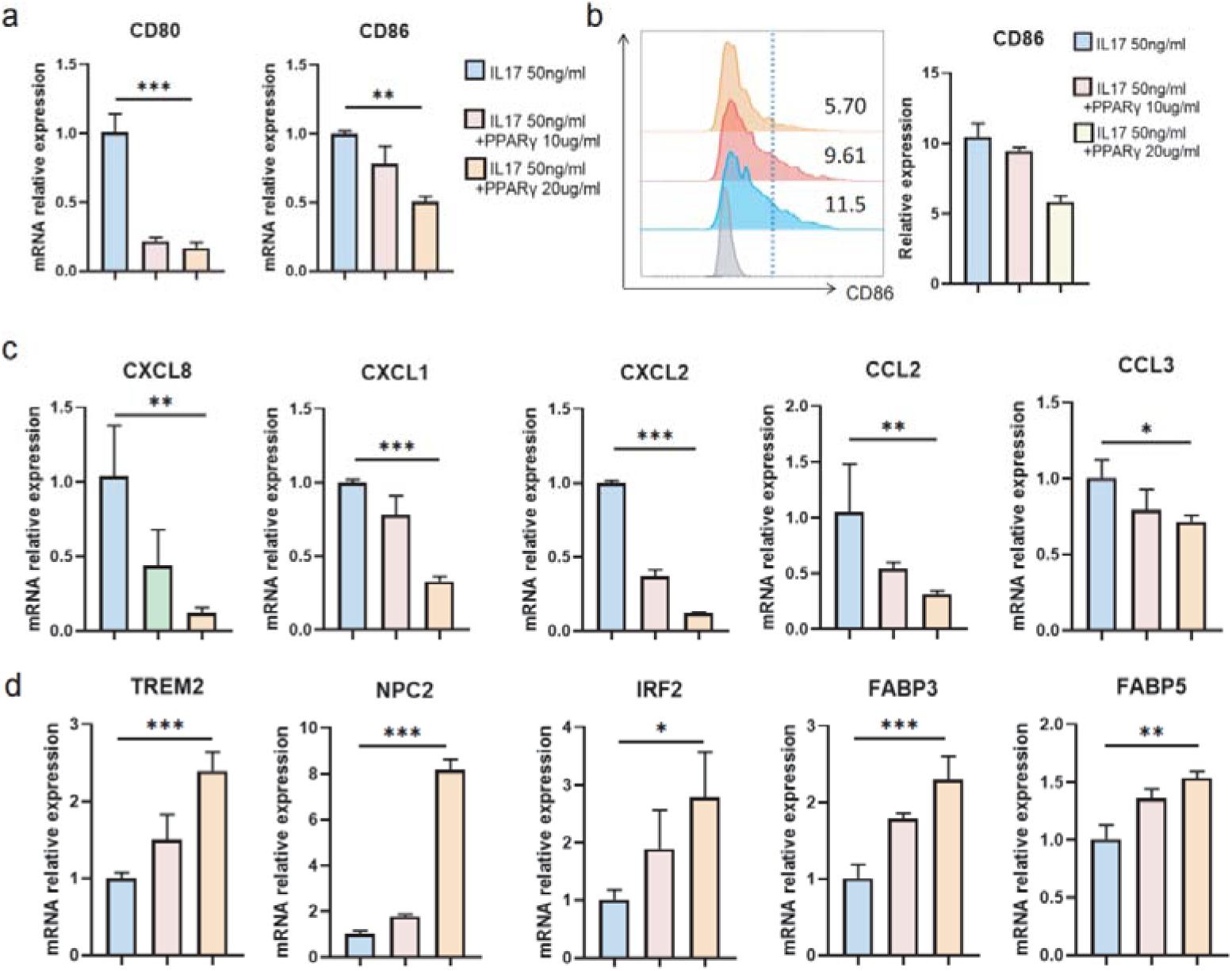
PPARγ Agonist Reverses Macrophage Disorganization Caused by IL-17A. **a,** mRNA expression levels of CD80, CD86 in macrophages after PPARγ agonist and IL-17A intervention. **b,** CD86 protein expression levels in macrophages after PPARγ agonist and IL-17A intervention. **c,** mRNA expression levels of CXCL2, CXCL3, CXCL8, CCL2, and CCL3 in macrophages after PPARγ agonist and IL-17A intervention. **d,** mRNA expression levels of TREM2,PLTP,APOE,FABP3 and FABP5 in macrophages after PPARγ agonist and IL-17A intervention. Significance indicated as *P < 0.05, **P < 0.01, ***P < 0.001, with ns denoting not significant. Statistical analysis was performed using the One-way ANOVA.

### Psoriasis Mouse Model Exacerbates Atherosclerosis, Ameliorated by IL-17A Inhibition

We designed in vivo experiments to verify the aggravating effect of psoriasis on atherosclerosis. Since subcutaneous injection is a trauma to mice that promotes psoriasis progression, we first explored the optimal concentration of IL-17A inhibitor. We intervened by injecting 100 μL of 10 μg/mL, 20 μg/mL, and 30 μg/mL secukinumab daily, every other day, and two days apart, respectively (Extended Data Fig7a). The results suggested that injecting 30 μg/mL secukinumab every other day was the most effective treatment for imiquimod-induced psoriasis in C57 mice (Extended Data Fig7b). Similarly, we found that alternate-day injections of 8 μg/ml ixekizumab effectively controlled psoriatic inflammation (Extended Data Fig8a,b).

In the formal experiment, we set up eight experimental groups: Con+PBS, Pso+PBS, As+PBS, Aspso+PBS, Con+IL-17 Inhibitor, Pso+IL-17 Inhibitor, As+IL-17 Inhibitor, and Aspso+IL-17 Inhibitor (six mice per group, the IL-17 inhibitor used was secukinumab). Peripheral blood serum, skin tissue, abdominal aorta, and extracted bone marrow progenitor cells were collected at the end of the experiment. The experimental flowchart is shown in Fig. 6a. In addition, we repeated the animal experiments for the As+IL-17 Inhibitor and Aspso+IL-17 Inhibitor groups using ixekizumab (six mice per group) to further validate the effect of IL-17 inhibition on comorbidity.

**Fig. 6.**
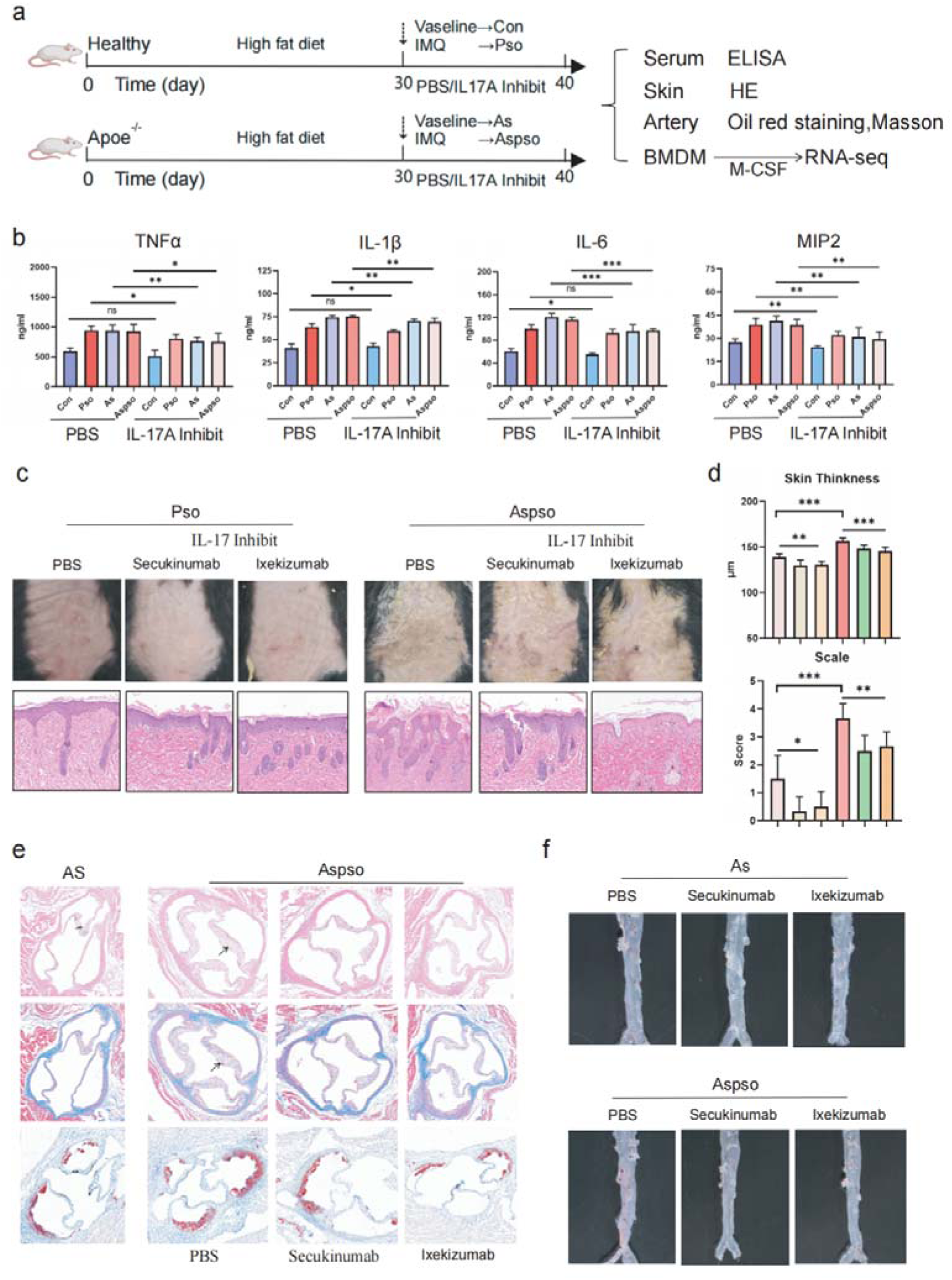
Psoriasis Mouse Model Aggravates the Severity of Atherosclerosis. **a,** Schematic diagram of the animal experiments. **b,** ELISA results showing the expression of TNF-α, IL-1β, IL-6, and MIP2 in the peripheral blood of each experimental group. **c,** Images of the backs of mice and corresponding hematoxylin and eosin (H&E) staining, demonstrating that atherosclerosis aggravates psoriasis. **d,** Changes in erythema and scaling scores in each experimental group. **e,** Hematoxylin and eosin (H&E) staining, Masson’s trichrome staining, and Oil Red O staining of the aortic valve. **f,** Images of mouse abdominal aortas stained with Oil Red O, showing that psoriasis aggravates atherosclerosis. Significance indicated as *P < 0.05, **P < 0.01, ***P < 0.001, with ns denoting not significant. Statistical analysis was performed using the t-test.

Peripheral blood serum was first collected for ELISA to detect relevant chemokines. Since CXCL8 is not expressed in mice, the comparable gene MIP2 was used as a substitute. ELISA showed that inflammatory chemokines such as TNF-α, IL-1β, IL-6, and MIP2 were significantly downregulated in the Pso, As, and Aspso groups after intervention with the IL-17A inhibitor (Fig. 6b). Comparing the changes in skin lesions among the Pso groups, Aspso groups, and corresponding IL-17A inhibitor intervention groups (Fig. 6c,d), the IL-17A inhibitor effectively ameliorated psoriasis inflammation. HE staining of skin lesions from all animal experiments is shown in Extended Data Fig9a, and immunofluorescence similarly suggested an increase in MIP2 in Aspso groups and improvement after IL-17 inhibitor intervention (Extended Data Fig9b).

As shown in Extended Data Fig10a, psoriasis induced by imiquimod aggravated plaque formation in the aortic arch, carotid artery, and aortic root of Apoe^-/-^ atherosclerotic mice. HE staining, Masson staining, and oil red O staining of the aortic valve revealed that imiquimod-induced psoriasis promoted valvular atherosclerotic plaque formation in Apoe^-/-^ mice, significantly reducing plaque collagen content and increasing lipid lesion area (Fig. 6e). Whole aortas were also examined by gross oil red O staining (Extended Data Fig10b-d).

As illustrated in Fig. 6f, oil red O staining showed that atherosclerosis was more severe in the abdominal aorta of Aspso groups, particularly in the segment corresponding to the dorsal skin area—where psoriasis was most pronounced.

Following IL-17A inhibitor treatment, the extent of atherosclerosis was significantly reduced.

### IL-17A Inhibitors Alleviate Atherosclerosis by Upregulating PPAR**_γ_**

We extracted bone marrow progenitor cells from the above mice and differentiated them into macrophages using M-CSF, followed by RNA-seq analysis. As shown in Fig. 7a, downregulation of PPARG (PPARγ) was more significant in the Aspso groups compared to the As groups. After IL-17A inhibitor intervention, the expression of PPARG (PPARγ) was upregulated to varying degrees. Meanwhile, inflammatory chemokines such as IL1B, IL12B, CCL3, CXCL1, and CXCL2 were significantly downregulated after IL-17A inhibitor intervention. Lipid metabolism-related genes such as TREM2, FABP3, and FABP5 were significantly downregulated in the Aspso groups and were reversed by IL-17A inhibitors.

**Fig. 7.**
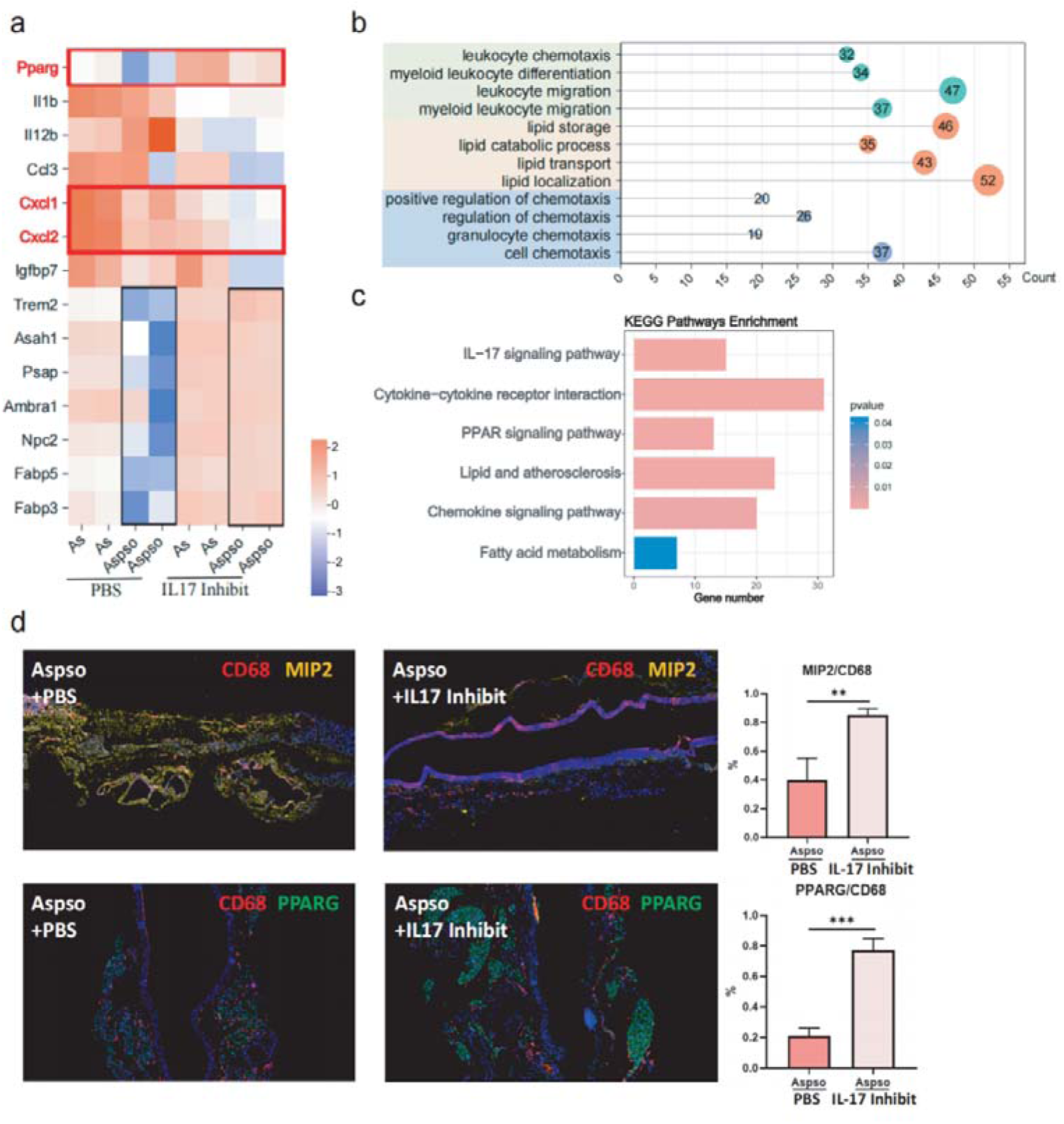
Inhibition of IL-17A Upregulates PPAR_γ_ Levels. **a,** RNA sequencing data showing mRNA changes in mouse bone marrow primary macrophages after IL-17A inhibition. **b,** GO enrichment analysis of differential genes in bone marrow primary macrophages between IL-17A inhibitor-treated and control groups. **c,** KEGG enrichment analysis of differential genes in bone marrow primary macrophages between IL-17A inhibitor-treated and control groups. **d,** Immunofluorescence showing changes in MIP2 and PPARγ in the abdominal aorta between IL-17A inhibitor-treated and control groups. Significance indicated as *P < 0.05, **P < 0.01, ***P < 0.001, with ns denoting not significant. Statistical analysis was performed using the t-test.

Differential genes of primary macrophages before and after IL-17A inhibitor intervention were subjected to enrichment analysis; GO enrichment analysis revealed differences in pathways of neutrophil chemotaxis, lipid metabolism, and chemokine secretion (Fig. 7b), and KEGG enrichment analysis revealed abnormal enrichment in the IL-17 pathway, PPAR signaling pathway, lipid metabolism and atherosclerosis pathway, and fatty acid and lipid metabolism pathway (Fig. 7c). Immunofluorescence confirmed the downregulation of MIP2 and upregulation of PPARγ after IL-17 inhibitor intervention (Fig. 7d). At the in vivo level, we verified that IL-17 interfered with macrophage-mediated progression of atherosclerosis by promoting inflammatory chemokine secretion and disrupting lipid metabolism.

In conclusion, psoriasis promotes the upregulation of pro-inflammatory factors such as CXCL2 and CXCL8 in macrophages through IL-17A-mediated downregulation of PPARγ. This regulation affects their pro-inflammatory interactions with endothelial cells and the immune microenvironment, and leads to macrophage dysfunction through the loss of sphingolipid metabolites via PPARγ, thereby mediating the destabilization of atherosclerotic plaques (Fig. 8).

**Fig. 8.**
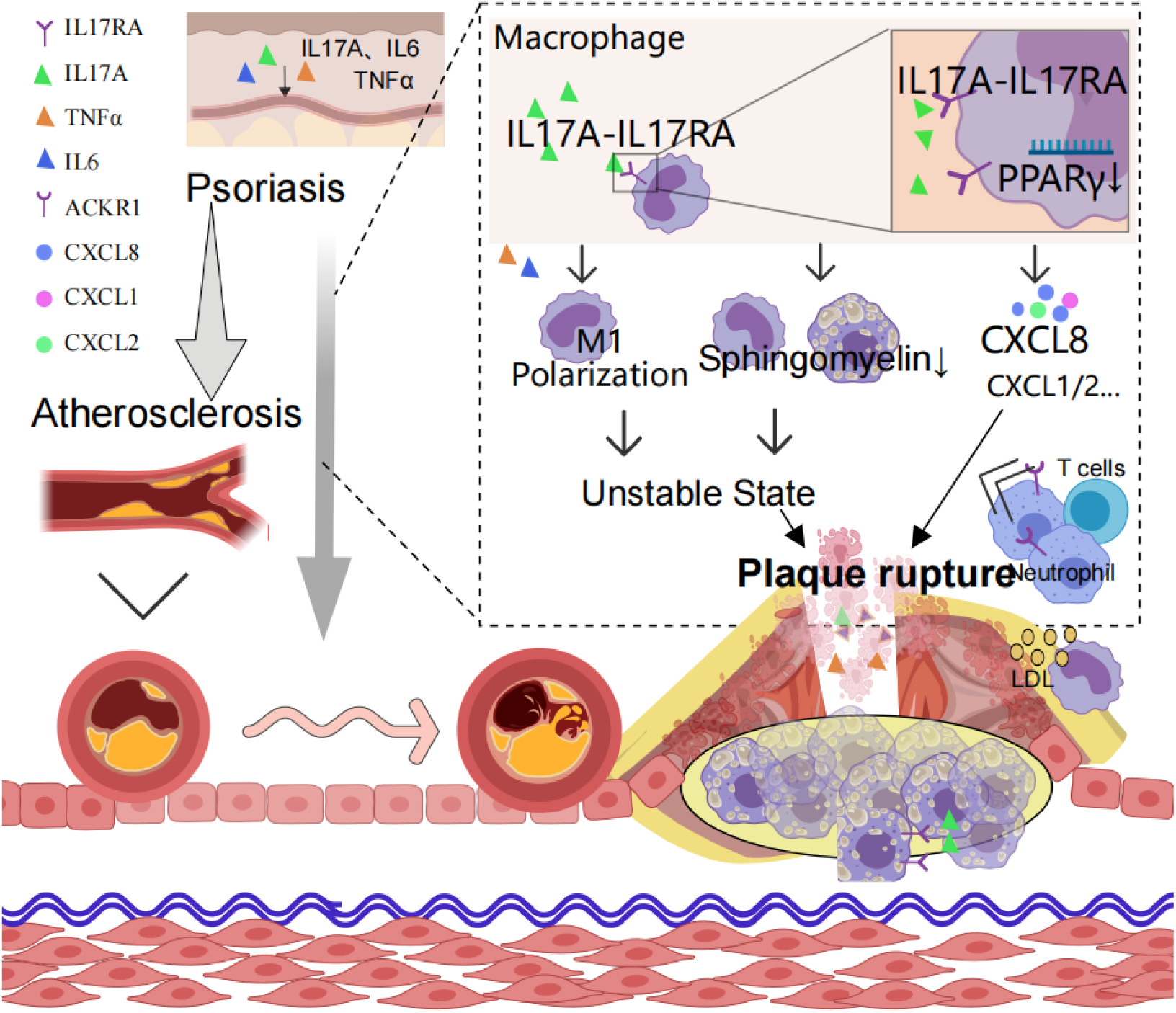
Mechanism Diagram. Psoriasis mediates atherosclerotic plaque destabilization through IL-17A–IL-17RA–mediated downregulation of PPARγ in macrophages, leading to macrophage disorganization characterized by increased inflammation, production of inflammatory factors, and downregulation of sphingomyelin components.

## DISCUSSION

Psoriasis and atherosclerosis share common pathological mechanisms and can exacerbate each other. In this study, we focused on macrophages as the core cells and proposed that the IL-17A/PPARγ axis is a mechanism by which psoriasis exacerbates atherosclerosis, leading to plaque instability. We suggest that timely use of IL-17A inhibitors can effectively reduce the aggravating effect of psoriasis on atherosclerosis.

Macrophages are key cells in the development of atherosclerosis, and IL-17A activates signaling pathways such as STAT3 by binding to macrophage surface receptors, promoting their polarization to the M1 type ^27, 28^. Meanwhile, IL-17A stimulates macrophages to secrete a large number of chemokines, such as CXCL1, CXCL8 (IL-8), and CCL2, exacerbating plaque instability ^29, 30^. In our study, single-cell transcriptome sequencing data revealed that chemokines CCL2, CCL3, CXCL2, CXCL3, and CXCL8 were upregulated in macrophages of psoriatic atherosclerosis plaques. Differential gene enrichment analysis between IL17RA_hi and IL-17RA_lo groups also showed enrichment of factors related to plaque progression and destabilization, such as chemokines and neutrophil recruitment. These findings were further confirmed by cellular experiments.

IL-17A, a key cytokine in the development and progression of psoriasis, is significantly upregulated in both peripheral blood and skin lesions of psoriasis patients ^24^. Its ability to promote the progression and destabilization of atherosclerotic plaques has been reported in previous literature. IL-17A promotes the secretion of various pro-inflammatory factors (e.g., IL-6, IL-8, GM-CSF) by macrophages and endothelial cells, attracting the migration of inflammatory cells such as neutrophils to the arterial wall and exacerbating local inflammatory responses ^30, 31^. In our study, samples from patients with both psoriasis and atherosclerosis were collected, and single-cell transcriptome sequencing of skin lesions and vascular tissues suggested that the receptor for IL-17A, IL-17RA, was upregulated. It was also demonstrated that IL-17RA was significantly upregulated in unstable plaques.

In addition to chemokines, the metabolic capabilities of macrophages also play a role in maintaining plaque stability. Phospholipids, as essential components of the cell membrane, enhance macrophage membrane fluidity and promote their phagocytic capacity ^32–35^. Previous studies have shown that psoriasis can impair lipid handling ^36^. In our study, phospholipid components such as sphingolipids and glycerophospholipids were significantly downregulated in macrophages after IL-17A treatment. Moreover, TREM2, a key gene regulating phospholipid metabolism, was also markedly downregulated following IL-17A intervention. IL-17A may impair phospholipid content in the membrane by disrupting sphingolipid metabolism, contributing to macrophage dysfunction and exacerbating the progression and instability of atherosclerotic plaques, although direct evidence is still lacking. Additionally, some studies suggest that IL-17A significantly enhances human macrophage macropinocytosis of modified LDL, driving foam cell formation, which may be associated with CD36, ABCA1, and ABCG1^37–39^, though no direct evidence has been established. The regulation of lipid metabolism in macrophages by IL-17A remains incompletely understood, representing a fertile area for future research.

PPARγ is a nuclear receptor encoded by the PPARG gene, mainly expressed in macrophages, and regulates cholesterol uptake, intracellular metabolism, and cytokine production and secretion. PPARγ has demonstrated protective effects in both psoriasis and atherosclerosis. In psoriasis, obesity induces downregulation of PPARγ in regulatory T cells (Tregs), leading to increased production of IL-17 ^40^. In atherosclerosis, PPARγ can slow disease progression by inhibiting the nuclear factor κB (NF-κB) pathway, reducing the expression of pro-inflammatory factors (e.g., IL-6, TNF-α), and inhibiting macrophage M1 polarization ^41^. Additionally, PPARγ can regulate lipid metabolism by modulating sphingolipid metabolism and increasing levels of anti-apoptotic molecules such as sphingosine-1-phosphate (S1P). S1P promotes macrophage survival and reduces foam cell death and necrotic core formation within plaques, thereby enhancing plaque stability ^42, 43^. PPARγ is also involved in the regulation of phospholipid metabolism, especially molecules like phosphatidylcholine and phosphatidylserine (PS). Phospholipids are important components of cell membranes, and their metabolic disorders affect membrane stability and macrophage function. PPARγ maintains membrane stability and functional integrity of macrophages by regulating the synthesis and degradation of these phospholipids ^44, 45^. In our study, the enrichment pathways related to lipid homeostasis, cytokine-mediated signaling pathway, myeloid leukocyte differentiation all involve PPARγ. Cellular experiments also confirmed that the chemokine and lipid metabolism disorders caused by IL-17A could be reversed by PPARγ agonists.

In addition to macrophages, we observed abnormalities in fibroblasts and endothelial cells in psoriatic atherosclerosis patients. Single-cell transcriptome sequencing revealed significant upregulation of the ACKR1 receptor in endothelial cells. Fibroblasts demonstrated increased expression of inflammatory factors. These findings suggest that psoriasis influences the atherosclerotic inflammatory immune microenvironment through IL-17A.

While psoriasis aggravates atherosclerosis, atherosclerosis likewise promotes psoriasis progression. It is recognized as a systemic inflammatory disease that can lead to upregulation of pro-inflammatory factors such as IL-1β, IL-6, and TNF-α in peripheral blood, which promote Th17 differentiation, IL-17 secretion, and psoriasis progression ^45^. We have previously found that MMP2, an important destabilizing factor of atherosclerotic plaques, can also promote the amplification of inflammation in psoriasis ^12, 13, 15^. Additionally, our study found that SGK1, which regulates macrophage inflammatory responses, foam cell formation, and cell survival in atherosclerosis, was downregulated in macrophages of psoriatic lesions, thereby promoting psoriasis inflammation ^46, 47^. These observations suggest the potential impact of atherosclerosis on psoriasis. In the present study, macrophages in skin lesions of psoriatic atherosclerosis patients demonstrated upregulation of chemokines such as CXCL8 along with IL-17RA upregulation. Endothelial cells and fibroblasts also exhibited higher pro-inflammatory properties. These findings suggest that IL-17A systemically affects both psoriasis and atherosclerosis.

In clinical practice, IL-17A inhibitors such as secukinumab and ixekizumab have been associated with cardiovascular protective effects while treating psoriasis ^48–50^. In our in vivo mouse experiments, we observed similar results. We constructed a co-morbid mouse model with both psoriasis and atherosclerosis, using mice with psoriasis or atherosclerosis alone as controls, and intervened with IL-17A inhibitors. The results revealed significant mutual exacerbation of psoriasis and atherosclerosis, consistent with previous findings. We found that the most significant aggravation of atherosclerosis occurred in the abdominal aorta corresponding to the back of the mice, possibly due to more pronounced inflammation in the psoriasis plaque area affecting vascular stability. Meanwhile, inhibition of IL-17A significantly improved both conditions. Peripheral blood ELISA demonstrated that IL-17A inhibitors downregulated pro-inflammatory factors such as IL-1β, IL-6, TNF-α, and MIP2, which is beneficial for both psoriasis and atherosclerosis. RNA-seq analysis of primary macrophages showed upregulation of PPARγ after IL-17A inhibitor intervention, demonstrating that these inhibitors systematically increase macrophage PPARγ expression and slow down the progression of both diseases.

However, this study has some limitations. First, in the in vivo validation of IL-17A promoting plaque destabilization, we only confirmed that psoriasis promotes the progression of atherosclerosis and alters plaque destabilization-associated genes expression, as an animal model of plaque destabilization is not yet available. Second, both psoriasis and atherosclerosis are long-term, chronic inflammatory diseases, and the model used in this study does not fully mimic the real-world phenomenon of these conditions mutually promoting each other throughout the disease course. Third, current randomized placebo-controlled clinical trials indicate that they exhibit neutral to beneficial effects on biomarkers associated with aortic vascular inflammation and cardiometabolic diseases ^53,54^. Finally, despite nearly a decade of clinical experience with secukinumab, the evidence supporting IL-17 monoclonal antibodies in reducing adverse cardiovascular events remains limited ^55^. This limitation may affect the direct translational potential of our findings.

In conclusion, this study revealed specific alterations in the immune microenvironment of atherosclerotic plaques due to psoriasis. Psoriasis downregulates PPARγ via the IL-17A/IL-17RA axis, resulting in the loss of macrophage sphingolipids and the production of large amounts of inflammatory factors (e.g., CXCL8), which mediate the progression of atherosclerosis and plaque destabilization. For patients with advanced atherosclerosis (unstable phase) and psoriasis, the early use of IL-17A inhibitors may help reduce psoriasis-related adverse cardiovascular outcomes.

## MATERIALS AND METHODS

### Skin and Atherosclerotic Plaque Samples

Patients with chronic plaque psoriasis diagnosed by a dermatologist and atherosclerosis diagnosed by a vascular surgeon were recruited for this study. Typical lesions were selected for biopsy in psoriasis patients, and carotid plaques were removed during surgery. All patients were free of autoimmune or systemic inflammatory diseases, and the lesion area and surrounding 5 cm had not received any therapeutic measures for at least two weeks before tissue removal. All participants provided written informed consent. The relevant patient data are presented in Extended Data Table1. Carotid plaque instability was determined through a comprehensive assessment combining high-resolution imaging modalities (ultrasound, MRI, and CTA) to evaluate plaque morphology and composition, clinical symptoms (history of TIA or stroke), and Masson staining. The study protocol was designed and conducted in accordance with the principles of the Declaration of Helsinki and was approved by the Ethical Review Committee of Huashan Hospital, Fudan University.

### Single-cell transcriptome sequencing

Carotid plaques and typical psoriatic skin lesions were collected from patients with both atherosclerosis and psoriasis. Libraries were sequenced on an Illumina NovaSeq 6000 to generate paired-end reads of 150 bp. Data processing, including quality control, read alignment (hg38), and gene quantification, was performed using Singleron GEXSCOPE software. Samples were then combined into individual expression matrices using the Singleron data integration pipeline. Atherosclerosis controls were obtained from the database GSE235436(51), and psoriasis lesion controls were obtained from self-assessed psoriasis lesion data.The spatial transcriptomics data were obtained from GSE241346(52). Patient details used for single-cell sequencing is provided in Extended Data Table2.

### Cell clustering and cell type annotation

Clustering analysis of the merged cell matrices was performed using the R package Seurat (v3.1.2). Low-quality cells with fewer than 500 transcripts or 100 genes, or with more than 10% mitochondrial gene expression, were filtered out. The expression level of each cell was normalized using the NormalizeData function with default parameters. The FindVariableFeatures function was applied to select highly variable genes. To integrate different samples, the Harmony function was used. The data were normalized and centered using the ScaleData function. Principal component analysis (PCA) was performed on the selected variable genes, and the top 20 principal components (PCs) were used for cell clustering and dimensionality reduction using UMAP. Cell clustering was performed using the FindNeighbors and FindClusters functions with the resolution parameter set to 1.0. Finally, cluster signature genes were identified using the FindAllMarkers function, and cell types were annotated based on these signature genes compared to known cell type markers.

### Enrichment analysis, cellular interactions, mimetic timing analysis

For Gene Ontology (GO) and Kyoto Encyclopedia of Genes and Genomes (KEGG) enrichment analyses, we used the R packages "enrichGO" and "enrichKEGG," respectively. These tools identify collections of genes significant in biological processes. Cellular interactions were analyzed using the R package "CellChat," which explores communication and signaling mechanisms between different cell types, revealing complex interactions. For dynamic changes during cell development, we used the R package Monocle2 to construct pseudotime trajectory analyses to show state transitions of cells over time and to capture potential developmental pathways and differentiation processes.

### Mice and treatment

C57BL/6J mice were purchased from Charles River Laboratories, Beijing, at 6–8 weeks of age. All mice were kept under specific pathogen-free conditions in the Department of Laboratory Animal Medicine of the university. To construct a mouse model of atherosclerosis, ApoE knockout mice on a C57BL/6 background were fed a 60% high-fat diet for one month, and the severity of atherosclerosis was subsequently detected using Oil Red O staining (Xavier, Wuhan, China). To induce moderate-to-severe psoriatic dermatitis, 62.5 mg of imiquimod (Mingxinlidi, Sichuan, China) was applied to a 2 cm × 2 cm area on the back, and an equal dose of petroleum jelly was used as a control. The IL-17A inhibitor, secukinumab (Novartis, Switzerland) and Ixekizumab (Lilly, the USA), are employed for the intervention. Ear thickness was measured using vernier calipers (Mitutoyo, Japan). Mice were scored for erythema and scaling with reference to (15). Animal experiments were performed in accordance with the guidelines in the university’s Guide for the Care and Use of Laboratory Animals.

### Cell culture

To extract primary macrophages from human peripheral blood, blood was collected from volunteer donors using anticoagulation tubes, and peripheral blood mononuclear cells (PBMCs) were isolated using density gradient centrifugation (e.g., Ficoll). The isolated PBMCs were suspended in RPMI 1640 medium containing 10% fetal bovine serum (FBS) and seeded into petri dishes. Monocytes were incubated at 37°C with 5% CO□ for 2 hours to promote adherence. Non-adherent cells were discarded and replaced with fresh complete medium containing macrophage colony-stimulating factor (M-CSF) at a concentration of 50 ng/mL. The culture was continued under the same conditions for 7 days, with the medium changed every 3 days, during which monocytes gradually differentiated into macrophages.

Similarly, to extract primary mononuclear macrophages from mice, femurs and tibiae were collected from C57BL/6 mice, and bone marrow cells were harvested by flushing with PBS. The extracted bone marrow cells were cultured in complete medium containing M-CSF (10 ng/mL) and incubated at 37°C with 5% CO□ for 3 days. Cells were collected after adherence was observed for subsequent experiments. Endothelial cells (Human Microvascular Endothelial Cell Line-1, HMEC-1,Procell Life Science & Technology Co., Ltd., Wuhan, China,Catalog #CL-0576) were cultured in RPMI 1640 medium supplemented with 10% fetal bovine serum (FBS; Gibco, Invitrogen, CA, USA).

### Reverse-transcription quantitative real-time polymerase change reaction

After the cultured cells were centrifuged, the pellet was collected and TRIzol reagent was added, followed by mixing through repeated aspiration. RNA was extracted according to the instructions of the RNA Extraction Kit (GeneStone Bio, China). The extracted RNA was subjected to reverse transcription using the Reverse Transcription Kit (Takara, China) to generate cDNA, which was stored at –20°C.

The expression levels of the relevant mRNAs were detected using a 7900HT Fast Real-Time PCR System (Thermo Fisher) with TaqMan Universal PCR Premix (Thermo Fisher #4304437) and TaqMan primers.Relevant primers in the text refer to Extended Data Table3.

### Immunohistochemistry

Slides were incubated with 3% H□O□ for 15 minutes to inhibit endogenous peroxidase activity. Sections were subjected to microwave antigen retrieval and incubated with primary antibodies at 4°C overnight. Antibodies used included IL-17RA(Abcam, Catalog #ab180904), CXCL8(HUABIO, Catalog #ER1706-67), PPARγ(Servicebio, Catalog #GB11164), and CD68(Servicebio, Catalog #GB115723). Sections were washed with PBS and incubated with goat anti-mouse IgG secondary antibody (Dako, Glostrup, Denmark) for 1 hour at room temperature. Afterwards, slides were washed again and treated with appropriate secondary antibodies, peroxidase (30 minutes), and diaminobenzidine (DAB) substrate, and finally imaged for analysis.

### Enzyme-linked immunosorbent assays

To detect the expression levels of TNF-α, IL-6, IL-1β, and MIP2 in mouse peripheral blood supernatants using ELISA, plasma or serum was first extracted from mouse blood (Hengyuan Biological Technology, China). ELISA plates were pre-coated with capture antibodies and incubated with blocking solution to prevent non-specific binding. Samples and standards were added and incubated under appropriate conditions. After incubation, the plates were washed several times with wash buffer, then HRP-labeled secondary antibodies were added and incubated. After washing, TMB substrate solution was added. After the color development reaction was terminated, absorbance was read at 450 nm using a microplate reader, and the concentration of each cytokine in the samples was calculated according to the standard curve.

### Flow cytometry

A total of 3 × 10□ cells were incubated with antibodies in PBS buffer containing 5% FBS and 0.1% sodium azide for 30 minutes on ice. Antibodies used included CD86 and CD163(Biolegend, USA). After incubation, cells were washed and resuspended in fixative (2% paraformaldehyde in PBS). Assays were performed using a BD FACSCalibur flow cytometer, and data were processed and analyzed using FlowJo 5.7.1 software.

### Liquid chromatography tandem mass spectrometry (LC-MS/MS)

Changes in protein expression in macrophages before and after IL-17A intervention were analyzed using LC-MS/MS. Cells were lysed, and proteins were extracted.

Proteins were digested with proteases, and peptides were separated by high-performance liquid chromatography (HPLC). Peptides were then subjected to mass spectrometry for mass-to-charge ratio determination and fragmentation analysis. Mass spectrometry data analysis software was used to identify and quantify the proteins, followed by functional analysis using bioinformatics tools.

### RNA sequencing

RNA was extracted using TRIzol reagent or RNA extraction kits (GeneStone Bio), and RNA concentration and integrity were assessed using NanoDrop and Agilent 2100 Bioanalyzer to ensure quality and purity. mRNA was enriched using poly-A tail enrichment or ribosomal RNA removal and subsequently fragmented. The mRNA was reverse transcribed to cDNA using reverse transcriptase, and libraries were constructed, including end repair, addition of A-tails, and adapter ligation. Libraries were amplified by PCR and checked for quality using Qubit and Agilent 2100. After quality control, high-throughput paired-end sequencing was performed using the Illumina NovaSeq 6000 platform, generating 50–100 million reads of 150 bp length for each sample. Raw data were quality-controlled using FastQC to remove low-quality reads and adapter-contaminated sequences, retaining clean reads. The clean reads were then aligned to the reference genome using alignment tools (STAR), and the number of reads or FPKM values for each gene was quantified using FeatureCounts or HTSeq.

### Immunofluorescence analysis

Tissues were fixed with 4% paraformaldehyde at room temperature and incubated with primary antibodies. Primary antibodies included mouse anti-CD68 (GB12192, Servicebio), mouse anti-PPARγ (sc-136250, Santa Cruz Biotechnology), and rabbit anti-MIP2 (GB11130, Servicebio). Cells were incubated overnight at 4°C with primary antibodies, followed by staining and incubation with secondary antibodies consisting of Alexa Fluor 555-labeled anti-rabbit IgG (1:2000, #4413, Cell Signaling Technology) and Alexa Fluor 488-labeled anti-mouse IgG (1:2000, #4408, Cell Signaling Technology) at room temperature. DAPI (Dako, Glostrup, Denmark) was used for nuclear counterstaining. Slides were analyzed using a confocal laser scanning microscope (FV-1000, Olympus, Tokyo, Japan).

### Statistics

Each experiment was performed with at least three biological replicates. Statistical analyses were performed using GraphPad Prism software (version 8.0, GraphPad Software, San Diego, USA) with unpaired two-tailed Student’s t-tests and One-way ANOVA. Results are presented as mean ± standard deviation (SD). Differences were considered statistically significant when the p-value was less than 0.05.

### Supplementary Material

Refer to Web version on PubMed Central for supplementary material.

## Acknowledgments

This study was funded by the National Natural Science Foundation of China (NSFC, grant number 82003382,52371249), the Bethune Charity Foundation (grant number J202301E036), the National Key Research and Development Program of China (2023YFC2508103), and the Clinical Research Plan of the Shanghai Health Development Center (grant numbers SHDC2020CR6022 and SHDC22022302).The studies involving human participants were reviewed and approved by the Ethics Committee of Huashan Hospital, Fudan University (2020-1202) and the Ethics Committee of Pudong Hospital, Fudan University (QWJWXK-24). The study was performed in accordance with the Declaration of Helsinki. All animal experiments were approved by the Ethics Committee of Huashan Hospital (approval no. 2020 JS-294).

## Data availability

All data relevant to this study are presented in the manuscript or supplementary materials. The scripts for the single-cell sequencing analysis can be accessed at https://github.com/Alex-0905-png/psoriatic-atherosclerosis

## References

1. W. H. Boehncke, M. P. Schön, Psoriasis. Lancet 386, 983–994 (2015).

2. Liu L, Cui S, Liu M, Huo X, Zhang G, Wang N. Psoriasis Increased the Risk of Adverse Cardiovascular Outcomes: A New Systematic Review and Meta-Analysis of Cohort Study. Front Cardiovasc Med 25;9:829709(2022).

3. H. Sharma, K. Mossman, R. C. Austin, Fatal attractions that trigger inflammation and drive atherosclerotic disease. Eur J Clin Invest 54, e14169 (2024).

4. M. J. Stiller, G. H. Pak, C. Kenny, L. Jondreau, I. Davis, S. Wachsman, J. L. Shupack. Elevation of fasting serum lipids in patients treated with low-dose cyclosporine for severe plaque-type psoriasis. An assessment of clinical significance when viewed as a risk factor for cardiovascular disease. J Am Acad Dermatol.27(3):434–8(1992).

5. H. Katz, J. Waalen, E. E. Leach.Acitretin in psoriasis: an overview of adverse effects. J Am Acad Dermatol. 41(3 Pt 2):S7–S12(1999).

6. G. R. Y. De Meyer, M. Zurek, P. Puylaert, W. Martinet, Programmed death of macrophages in atherosclerosis: mechanisms and therapeutic targets. Nat Rev Cardiol 21, 312–325 (2024).

7. O. J. Waring, N. T. Skenteris, E. A. L. Biessen, M. Donners, Two-faced Janus: the dual role of macrophages in atherosclerotic calcification. Cardiovasc Res 118, 2768–2777 (2022).

8. C. Weber, A. J. R. Habenicht, P. von Hundelshausen, Novel mechanisms and therapeutic targets in atherosclerosis: inflammation and beyond. Eur Heart J 44, 2672–2681 (2023).

9. U. M. Breland, B. Halvorsen, J. Hol, E. Øie, G. P. Berne, A. Yndestad, C. Smith, K. Otterdal, U. Hedin, T. Waehre, W. J. Sandberg, S. S. Frøland, G. Haraldsen, L. Gullestad, J. K. Damås, G. K. Hansson, P. Aukrust, A potential role of the CXC chemokine GROalpha in atherosclerosis and plaque destabilization: downregulatory effects of statins. Arterioscler Thromb Vasc Biol 28, 1005–1011 (2008).

10. O. Soehnlein, M. Drechsler, Y. Döring, D. Lievens, H. Hartwig, K. Kemmerich, A. O. Gómez, M. Mandl, S. Vijayan, D. Projahn, C. D. Garlichs, R. R. Koenen, M. Hristov, E. Lutgens, A. Zernecke, C. Weber, Distinct functions of chemokine receptor axes in the atherogenic mobilization and recruitment of classical monocytes. EMBO Mol Med 5, 471–481 (2013).

11. D. A. Chistiakov, A. N. Orekhov, Y. V. Bobryshev, Chemokines and Relevant microRNAs in the Atherogenic Process. Mini Rev Med Chem 18, 597–608 (2018).

12. M. Kuzuya, K. Nakamura, T. Sasaki, X. W. Cheng, S. Itohara, A. Iguchi, Effect of MMP-2 deficiency on atherosclerotic lesion formation in apoE-deficient mice. Arterioscler Thromb Vasc Biol 26, 1120–1125 (2006).

13. S. Momi, E. Falcinelli, E. Petito, G. C. Taranta, A. Ossoli, P. Gresele, Matrix metalloproteinase-2 on activated platelets triggers endothelial PAR-1 initiating atherosclerosis. Eur Heart J 43, 504–514 (2022).

14. P. K. Shah, E. Falk, J. J. Badimon, A. F. Ortiz, A. Mailhac, G. V. Levy, J. T. Fallon, J. Regnstrom, V. Fuster, Human monocyte-derived macrophages induce collagen breakdown in fibrous caps of atherosclerotic plaques. Potential role of matrix-degrading metalloproteinases and implications for plaque rupture. Circulation 92, 1565–1569 (1995).

15. C. Dong, J. Lin, X. Lu, J. Zhu, L. Lin, J. Xu, J. Du, Fibroblasts with high matrix metalloproteinase 2 expression regulate CD8+ T-cell residency and inflammation via CD100 in psoriasis. Br J Dermatol 191, 405–418 (2024).

16. R. Kodali, M. Hajjou, A. B. Berman, M. B. Bansal, S. Zhang, J. J. Pan, A. D. Schecter, Chemokines induce matrix metalloproteinase-2 through activation of epidermal growth factor receptor in arterial smooth muscle cells. Cardiovasc Res 69, 706–715 (2006).

17. C. Diaz-Canestro, Y. M. Puspitasari, L. Liberale, T. J. Guzik, A. J. Flammer, N. R. Bonetti, P. Wüst, S. Costantino, F. Paneni, A. Akhmedov, Z. Varga, S. Ministrini, J. H. Beer, F. Ruschitzka, M. Hermann, T. F. Lüscher, I. Sudano, G. G. Camici, MMP-2 knockdown blunts age-dependent carotid stiffness by decreasing elastin degradation and augmenting eNOS activation. Cardiovasc Res 118, 2385–2396 (2022).

18. N. Samah, A. Ugusman, A. A. Hamid, N. Sulaiman, A. Aminuddin, Role of Matrix Metalloproteinase-2 in the Development of Atherosclerosis among Patients with Coronary Artery Disease. Mediators Inflamm 2023, 9715114 (2023).

19. S. V. Subas, V. Mishra, V. Busa, I. Antony, S. Marudhai, M. Patel, I. Cancarevic, Cardiovascular Involvement in Psoriasis, Diagnosing Subclinical Atherosclerosis, Effects of Biological and Non-Biological Therapy: A Literature Review. Cureus 12, e11173 (2020).

20. Y. Wang, J. Zang, C. Liu, Z. Yan, D. Shi, Interleukin-17 Links Inflammatory Cross-Talks Between Comorbid Psoriasis and Atherosclerosis. Front Immunol 13, 835671 (2022).

21. S. Kotla, N. K. Singh, M. R. Heckle, G. J. Tigyi, G. N. Rao, The transcription factor CREB enhances interleukin-17A production and inflammation in a mouse model of atherosclerosis. Sci Signal 6, ra83 (2013).

22. S. Chen, K. Shimada, W. Zhang, G. Huang, T. R. Crother, M. Arditi, IL-17A is proatherogenic in high-fat diet-induced and Chlamydia pneumoniae infection-accelerated atherosclerosis in mice. J Immunol 185, 5619–5627 (2010).

23. J. Wang, Y. Gao, Y. Yuan, H. Wang, Z. Wang, X. Zhang, Th17 Cells and IL-17A in Ischemic Stroke. Mol Neurobiol 61, 2411–2429 (2024).

24. S. Bozinovski, H. J. Seow, S. P. J. Chan, D. Anthony, J. McQualter, M. Hansen, B. J. Jenkins, G. P. Anderson, R. Vlahos, Innate cellular sources of interleukin-17A regulate macrophage accumulation in cigarette- smoke-induced lung inflammation in mice. Clin Sci (Lond) 129, 785–796 (2015).

25. S. Fukuzaki, R. F. Righetti, T. M. D. Santos, L. d. N. Camargo, L. R. C. R. B. Aristóteles, F. C. R. Souza, A. C. Garrido, B. M. Saraiva-Romanholo, E. A. Leick, C. M. Prado, M. d. A. Martins, I. d. F. L. C. Tibério, Preventive and therapeutic effect of anti-IL-17 in an experimental model of elastase-induced lung injury in C57Bl6 mice. Am J Physiol Cell Physiol 320, C341–c354 (2021).

26. S. L. Gaffen, Structure and signalling in the IL-17 receptor family. Nat Rev Immunol 9, 556–567 (2009).

27. I. Tabas, K. E. Bornfeldt, Macrophage Phenotype and Function in Different Stages of Atherosclerosis. Circ Res 118, 653–667 (2016).

28. J. Nordlohne, S. von Vietinghoff, Interleukin 17A in atherosclerosis - Regulation and pathophysiologic effector function. Cytokine 122, 154089 (2019).

29. M. Akhavanpoor, H. Akhavanpoor, C. A. Gleissner, S. Wangler, A. O. Doesch, H. A. Katus, C. Erbel, The Two Faces of Interleukin-17A in Atherosclerosis. Curr Drug Targets 18, 863–873 (2017).

30. S. Taleb, A. Tedgui, Z. Mallat, IL-17 and Th17 cells in atherosclerosis: subtle and contextual roles. Arterioscler Thromb Vasc Biol 35, 258–264 (2015).

31. G. Allam, A. Abdel-Moneim, A. M. Gaber, The pleiotropic role of interleukin-17 in atherosclerosis. Biomed Pharmacother 106, 1412–1418 (2018).

32. A. Weigert, N. Weis, B. Brüne, Regulation of macrophage function by sphingosine-1-phosphate. Immunobiology 214, 748–760 (2009).

33. D. Schlam, J. Canton, M. Carreño, H. Kopinski, S. A. Freeman, S. Grinstein, G.D. Fairn, Gliotoxin Suppresses Macrophage Immune Function by Subverting Phosphatidylinositol 3,4,5-Trisphosphate Homeostasis. mBio 7, e02242 (2016).

34. J. A. Wali, N. Jarzebska, D. Raubenheimer, S. J. Simpson, R. N. Rodionov, J. F. O’Sullivan, Cardio-Metabolic Effects of High-Fat Diets and Their Underlying Mechanisms-A Narrative Review. Nutrients 12 (2020).

35. Z. Lu, Y. Li, W. Syn, Z. Wang, M. F. Lopes-Virella, T. J. Lyons, Y. Huang, Amitriptyline inhibits nonalcoholic steatohepatitis and atherosclerosis induced by high-fat diet and LPS through modulation of sphingolipid metabolism. Am J Physiol Endocrinol Metab 318, E131–e144 (2020).

36. M. Holzer, P. Wolf, S. Curcic, R. Birner-Gruenberger, W. Weger, M. Inzinger, D. El-Gamal, C. Wadsack, A. Heinemann, G. Marsche, Psoriasis alters HDL composition and cholesterol efflux capacity. J Lipid Res 53, 1618–1624 (2012).

37. D. R. Michael, T. G. Ashlin, C. S. Davies, H. Gallagher, T. W. Stoneman, M. L. Buckley, D. P. Ramji, Differential regulation of macropinocytosis in macrophages by cytokines: implications for foam cell formation and atherosclerosis. Cytokine 64, 357–361 (2013).

38. A. Owczarczyk-Saczonek, W. Placek, Interleukin-17 as a factor linking the pathogenesis of psoriasis with metabolic disorders. Int J Dermatol 56, 260–268 (2017).

39. T. R. Cimato, B. A. Palka, J. K. Lang, R. F. Young, LDL cholesterol modulates human CD34+ HSPCs through effects on proliferation and the IL-17 G-CSF axis. PLoS One 8, e73861 (2013).

40. P. Sivasami, C. Elkins, P. P. Diaz-Saldana, K. Goss, A. Peng, M. H. 4th, J. Bae, M. Xu, B. P. Pollack, E. M. Horwitz, C. D. Scharer, L. Seldin, C. Li, Obesity-induced dysregulation of skin-resident PPARγ(+) Treg cells promotes IL-17A-mediated psoriatic inflammation. Immunity 56, 1844–1861.e1846 (2023).

41. Y. Pu, C. K. Cheng, H. Zhang, J. Luo, L. Wang, B. Tomlinson, Y. Huang, Molecular mechanisms and therapeutic perspectives of peroxisome proliferator-activated receptor α agonists in cardiovascular health and disease. Med Res Rev 43, 2086–2114 (2023).

42. A. L. Catapano, A. Pirillo, F. Bonacina, G. D. Norata, HDL in innate and adaptive immunity. Cardiovasc Res 103, 372–383 (2014).

43. O. L. Manzo, J. Nour, L. Sasset, A. Marino, L. Rubinelli, S. Palikhe, M. Smimmo, Y. Hu, M. R. Bucci, A. Borczuk, O. Elemento, J. K. Freed, G. D. Norata, A. D. Lorenzo, Rewiring Endothelial Sphingolipid Metabolism to Favor S1P Over Ceramide Protects From Coronary Atherosclerosis. Circ Res 134, 990–1005 (2024).

44. S. J. Bensinger, P. Tontonoz, Integration of metabolism and inflammation by lipid-activated nuclear receptors. Nature 454, 470–477 (2008).

45. Y. Li, Y. Pan, X. Zhao, S. Wu, F. Li, Y. Wang, B. Liu, Y. Zhang, X. Gao, Y. Wang, H. Zhou, Peroxisome proliferator-activated receptors: A key link between lipid metabolism and cancer progression. Clin Nutr 43, 332–345 (2024).

46. C. Dong, J. Lin, Y. Wang, J. Zhu, L. Lin, J. Xu, J. Du, Exploring the Common Pathogenic Mechanisms of Psoriasis and Atopic Dermatitis: The Interaction between SGK1 and TIGIT Signaling Pathways. Inflammation 10.1007/s10753-024-02115-1 (2024).

47. O. Borst, M. Schaub, B. Walker, E. Schmid, P. Münzer, J. Voelkl, I. Alesutan, J. M. Rodríguez, S. Vogel, T. Schoenberger, K. Metzger, D. Rath, A. Umbach, D. Kuhl, I. I. Müller, P. Seizer, T. Geisler, M. Gawaz, F. Lang, Pivotal role of serum- and glucocorticoid-inducible kinase 1 in vascular inflammation and atherogenesis. Arterioscler Thromb Vasc Biol 35, 547–557 (2015).

48. Y. A. Elnabawi, A. K. Dey, A. Goyal, J. W. Groenendyk, J. H. Chung, A. D. Belur, J. Rodante, C. L. Harrington, H. L. Teague, Y. Baumer, A. Keel, M. P. Playford, V. Sandfort, M. Y. Chen, B. Lockshin, J. M. Gelfand, D. A. Bluemke, N. N. Mehta, Coronary artery plaque characteristics and treatment with biologic therapy in severe psoriasis: results from a prospective observational study. Cardiovasc Res 115, 721–728 (2019).

49. H. Choi, D. E. Uceda A. K. Dey, K. M. Abdelrahman, M. Aksentijevich, J. A. Rodante, Y. A. Elnabawi, A. Reddy, A. Keel, J. Erb-Alvarez, H. Teague, M. P. Playford, W. Zhou, M. Y. Chen, J. M. Gelfand, D. A. Bluemke, A. Buckler, N. N. Mehta, Treatment of Psoriasis With Biologic Therapy Is Associated With Improvement of Coronary Artery Plaque Lipid-Rich Necrotic Core: Results From a Prospective, Observational Study. Circ Cardiovasc Imaging 13, e011199 (2020).

50. Y. A. Elnabawi, E. K. Oikonomou, A. K. Dey, J. Mancio, J. A. Rodante, M. Aksentijevich, H. Choi, A. Keel, J. Erb-Alvarez, H. L. Teague, A. A. Joshi, M. P. Playford, B. Lockshin, A. D. Choi, J. M. Gelfand, M. Y. Chen, D. A. Bluemke, C. Shirodaria, C. Antoniades, N. N. Mehta, Association of Biologic Therapy With Coronary Inflammation in Patients With Psoriasis as Assessed by Perivascular Fat Attenuation Index. JAMA Cardiol 4, 885–891 (2019).

51. N. Eberhardt, M. G. Noval, R. Kaur, L. Amadori, M. Gildea, S. Sajja, D. Das, B. Cilhoroz, O. Stewart, D. M. Fernandez, R. Shamailova, A. V. Guillen, S. Jangra, M. Schotsaert, J. D. Newman, P. Faries, T. Maldonado, C. Rockman, A. Rapkiewicz, K. A. Stapleford, N. Narula, K. J. Moore, C. Giannarelli, SARS-CoV-2 infection triggers pro-atherogenic inflammatory responses in human coronary vessels. Nat Cardiovasc Res 2(10):899–916 (2023).

52. K. Theofilatos, S. Stojkovic, M. Hasman, S. W. v. d. Laan, F. Baig, J. Barallobre-Barreiro, L. E. Schmidt, S. Yin, X. Yin, S. Burnap, B. Singh, J. Popham, O. Harkot, S. Kampf, M. C. Nackenhorst, A. Strassl, C. Loewe, S. Demyanets, C. Neumayer, M. Bilban, C. Hengstenberg, K. Huber, G. Pasterkamp, J. Wojta, M. Mayr, Proteomic Atlas of Atherosclerosis: The Contribution of Proteoglycans to Sex Differences, Plaque Phenotypes, and Outcomes. Circ Res 133(7):542–558(2023).

53. J. M. Gelfand, D. B. Shin, K. C. Duffin, A. W. Armstrong, A. Blauvelt, S. K. Tyring, A. Menter, S. Gottlieb, B. N. Lockshin, E. L. Simpson, F. Kianifard, R. P. Sarkar, E. Muscianisi, J. Steadman, M. A. Ahlman, M. P. Playford, A. A. Joshi, A. K. Dey, T. J. Werner, A. Alavi, N. N. Mehta.A Randomized Placebo-Controlled Trial of Secukinumab on Aortic Vascular Inflammation in Moderate-to-Severe Plaque Psoriasis (VIP-S).J Invest Dermatol 140(9):1784–1793.e2(2020).

54. E. v. Stebut, K. Reich, D. Thaçi, W. Koenig, A. Pinter, A. Körber, T. Rassaf, A. Waisman, V. Mani, D. Yates, J. Frueh, C. Sieder, N. Melzer, N. N. Mehta, T. Gori. Impact of Secukinumab on Endothelial Dysfunction and Other Cardiovascular Disease Parameters in Psoriasis Patients over 52 Weeks. J Invest Dermatol 139(5):1054–1062(2019).

55. L. P. Vegas, P. L. Corvoisier, L. Penso, M. Paul, E. Sbidian, P. Claudepierre.Risk of major adverse cardiovascular events in patients initiating biologics/apremilast for psoriatic arthritis: a nationwide cohort study. Rheumatology (Oxford) 11;61(4):1589–1599(2022).

