## Supplemental Figures for "IL-17A-driven Macrophage Disorder Promotes Plaque Instability in Psoriatic Atherosclerosis"

**Supplemental Material**

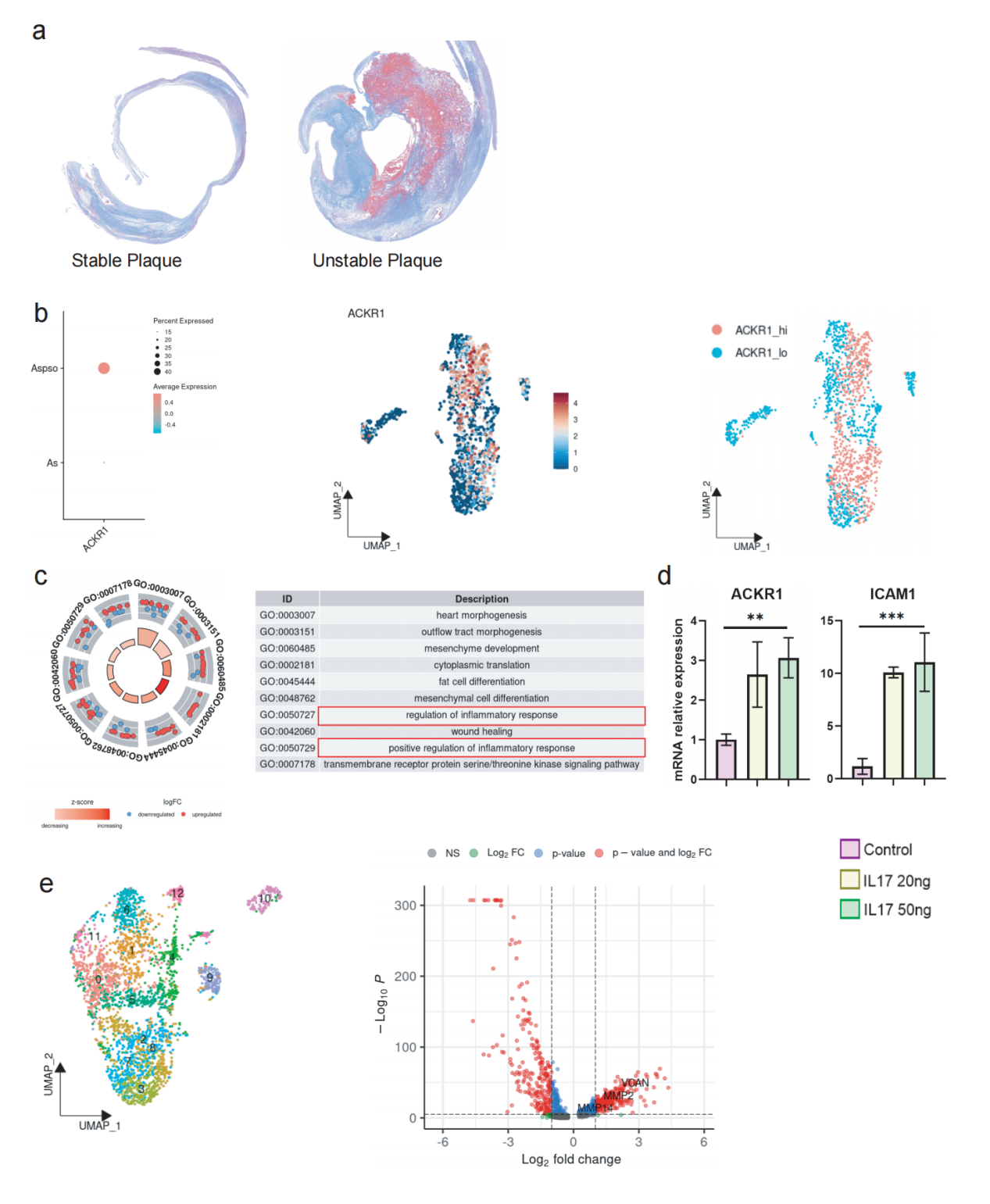

**Extended Data Fig1. Enhanced Inflammation of Endothelial Cells and Fibroblasts in Co-Morbid Plaques**

**a,** Masson staining of stable and unstable plaques;

**b,** Upregulation of ACKR1 expression in endothelial cells in psoriatic atherosclerosis plaques;

**c,** GO enrichment analysis of differential genes in endothelial cells with high and low ACKR1 expression;

**d,** Expression levels of ACKR1 and ICAM1 mRNAs in endothelial cells after intervention with CXCL8 recombinant protein;

**e,** Volcano plot illustrates the differentially expressed genes between fibroblasts associated with comorbidities and those exclusively from plaques.

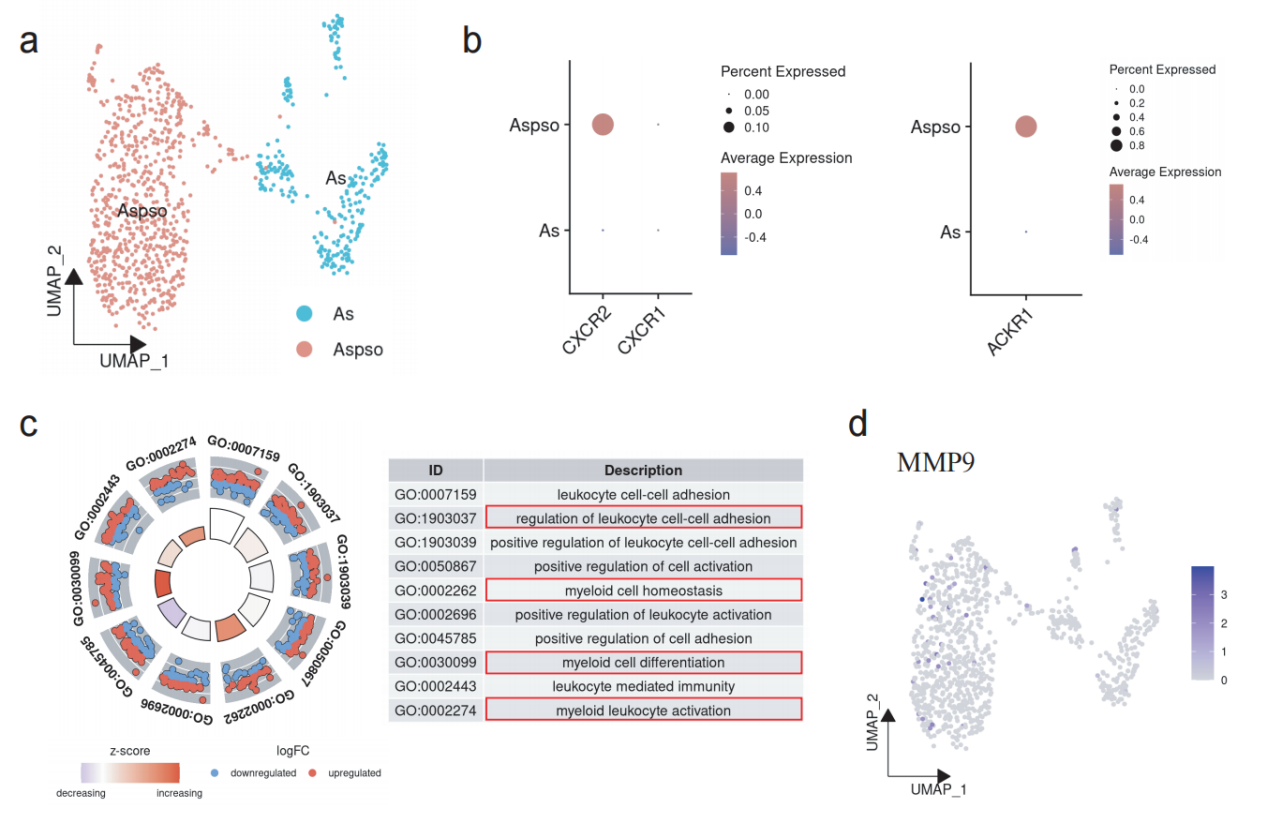

**Extended Data Fig2.Upregulation of CXCL8 Receptors in Myeloid Cells of Comorbid Plaques**

**a,** UMAP plot displaying the tissue origins of myeloid cells;

**b,** Dot plot showing the expression levels of CXCL8 receptors CXCR1/2 and ACKR1 in the Aspso and As groups;

**c,** Gene Ontology (GO) enrichment analysis of differentially expressed genes between the Aspso and As groups;

**d,** Expression levels of MMP9 in myeloid cells of the Aspso and As groups.

Aspso: Patients with both psoriasis and carotid plaques; As: Patients with carotid plaques only.

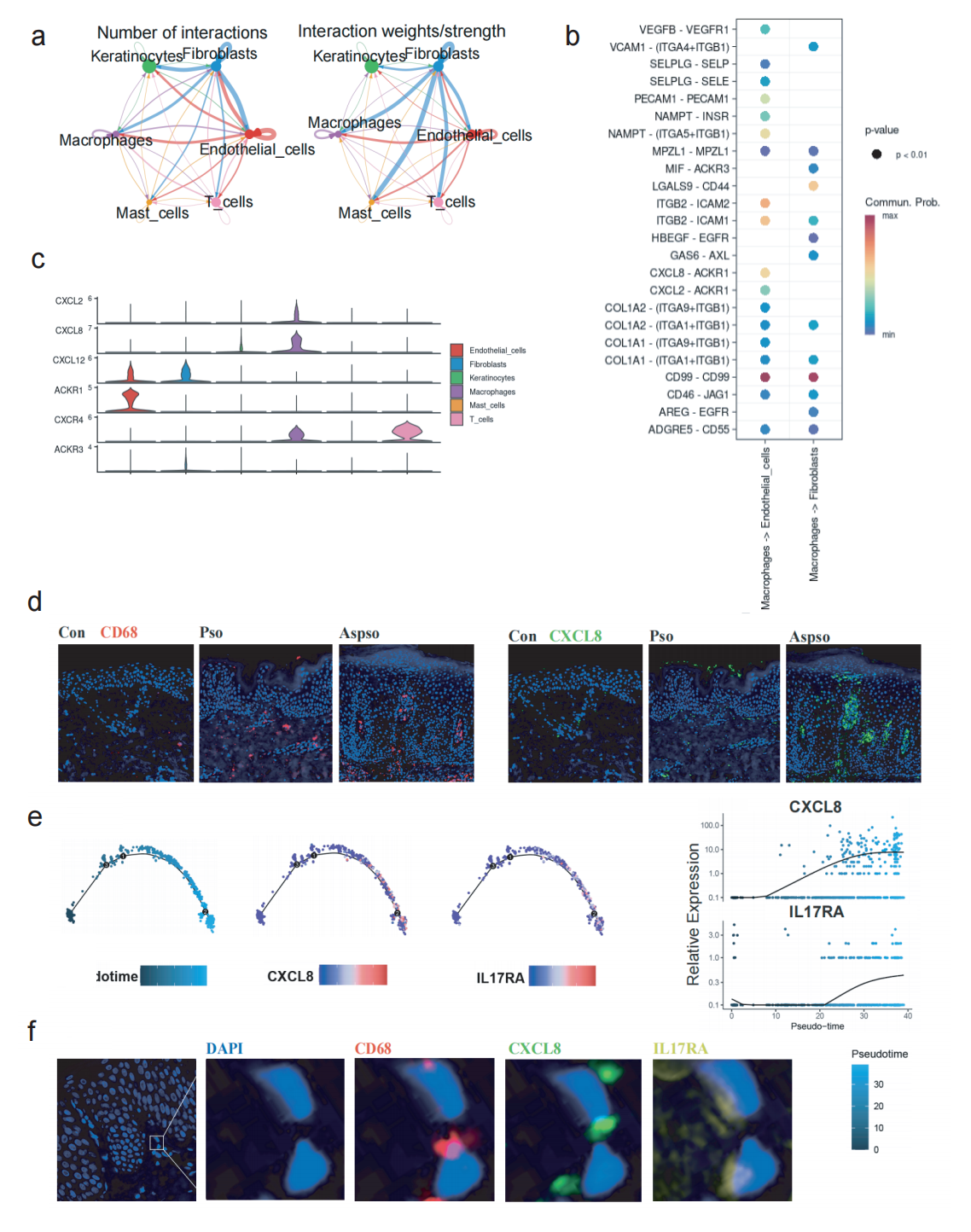

**Extended Data Fig3.** **Upregulation of Macrophage Chemokines in Skin Lesions Accompanied by IL-17RA Alterations**

**a,** Number of interactions and interaction weights among various cell types in co-morbid skin lesions;

**b,** Receptor-ligand interactions between macrophages and endothelial cells and fibroblasts in co-morbid skin lesions;

**c,** Expression levels of CXCL signaling molecules in various cell types within co-morbid lesions;

**d,** Immunofluorescence showing CXCL8 expression in co-morbid psoriatic lesions and psoriasis-only lesions;

**e,** Pseudotime analysis showing that IL-17RA upregulation is accompanied by CXCL8 upregulation;

**f,** Immunofluorescence showing co-localization of IL-17RA and CXCL8.

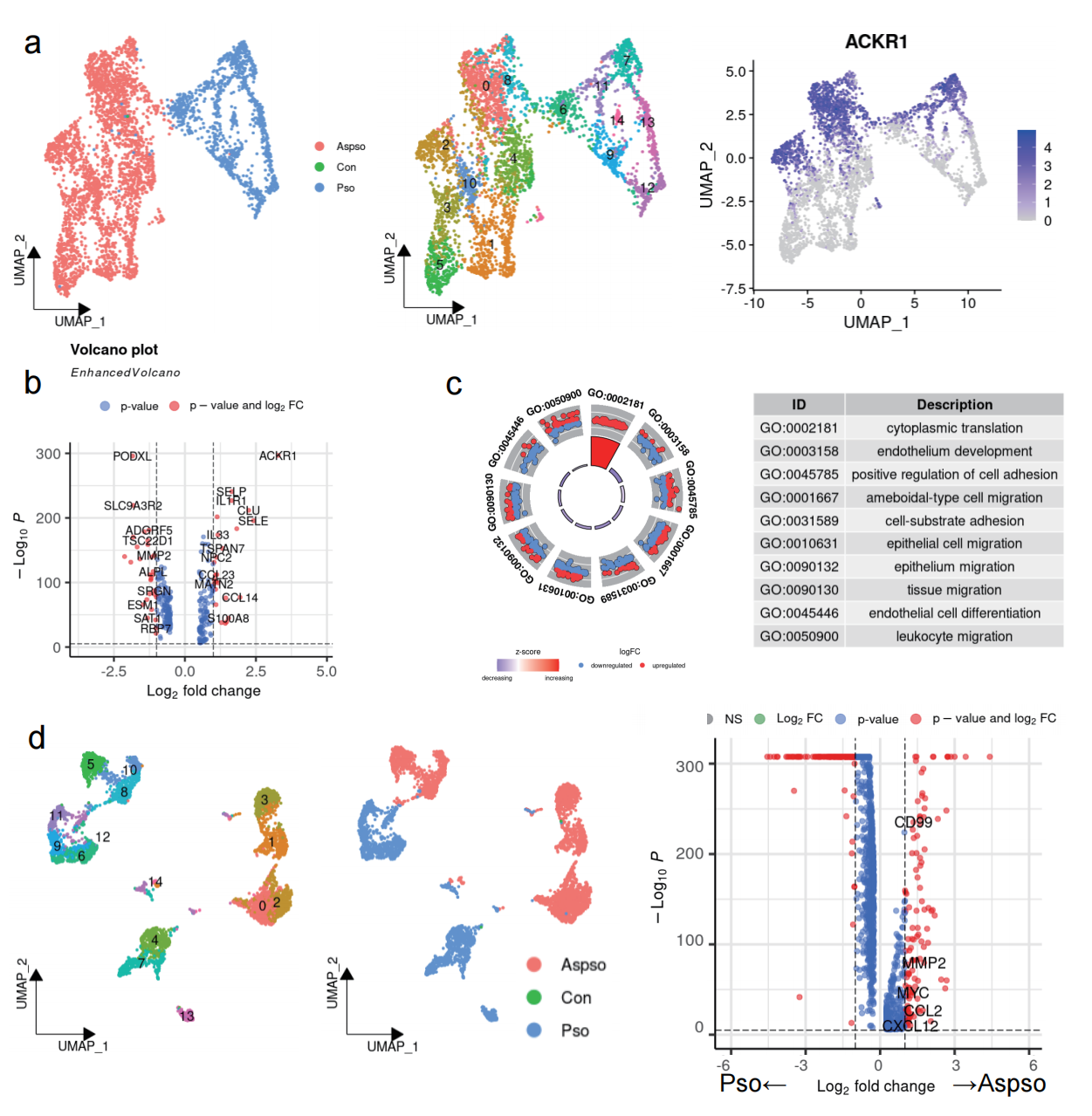

**Extended Data Fig4. Enhanced Inflammation of Endothelial Cells and Fibroblasts in Skin Lesions**

**a,** Upregulation of endothelial cell ACKR1 expression in co-morbid skin lesions;

**b,** Volcano plot showing differential genes in endothelial cells with high and low ACKR1 expression;

**c,** GO enrichment analysis of differential genes in endothelial cells with high and low ACKR1 expression;

**d,** Differential gene expression in fibroblasts from co-morbid lesions compared to psoriasis-only lesions.

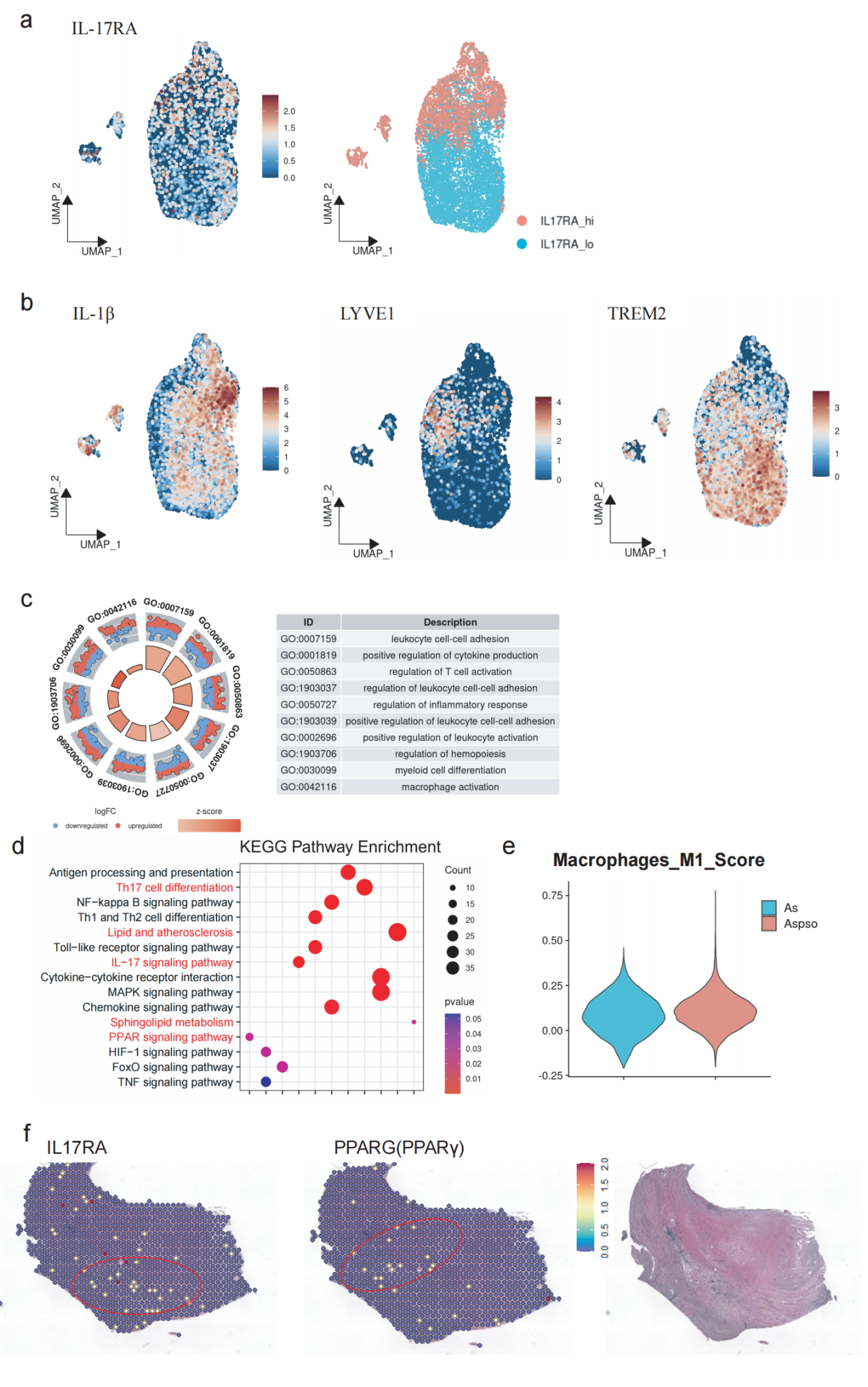

**Extended Data Fig5. Downregulation of PPARγ Expression in Regions of High IL-17RA Expression in Plaques**

**a,** Macrophages were categorized into two subpopulations, IL17RA_hi and IL17RA_lo, based on their IL17RA expression levels;

**b,** UMAP visualization of the expression distribution of IL-1β, LVYE1, and TREM2 in macrophages;

**c,** Gene Ontology (GO) enrichment pathways of differentially expressed genes between the Aspso and As groups;

**d,** Kyoto Encyclopedia of Genes and Genomes (KEGG) enrichment analysis of differentially expressed genes between the Aspso and As groups;

**e,** M1 polarization gene set scores in macrophages of the Aspso and As groups;

**f,** Single-cell sequencing combined with spatial transcriptomics showing that upregulated sites of IL-17RA in plaques are accompanied by downregulation of PPARγ.

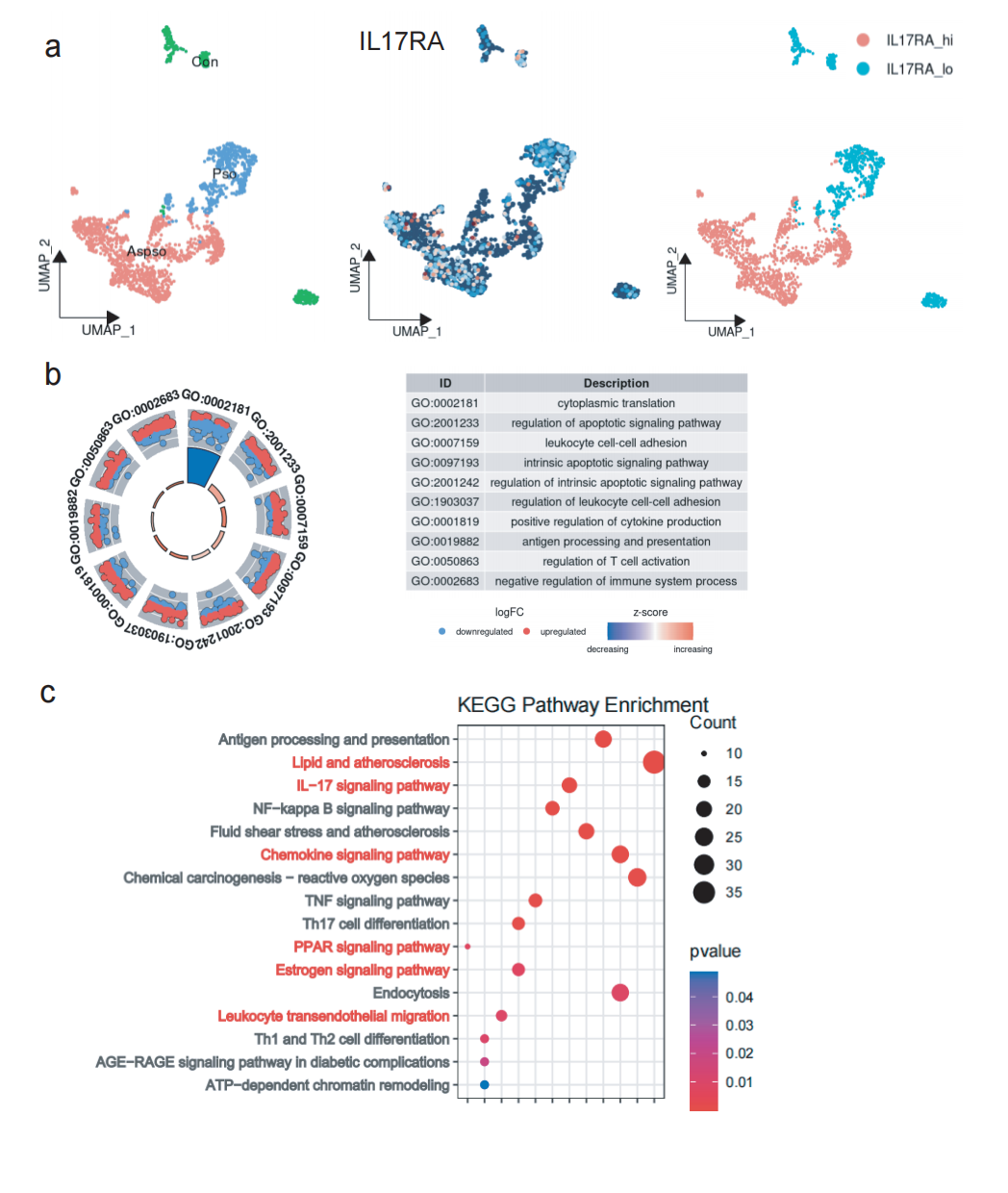

**Extended Data Fig6. Increased Macrophage Inflammation in Co-Morbid Psoriatic Lesions**

**a,** UMAP demonstrating tissue origin in co-morbid psoriatic lesions;

**b,** GO enrichment analysis of differential genes between macrophages in co-morbid psoriatic lesions and psoriasis-only lesions;

**c,** KEGG enrichment analysis of differential genes between macrophages in co-morbid psoriatic lesions and psoriasis-only lesions.

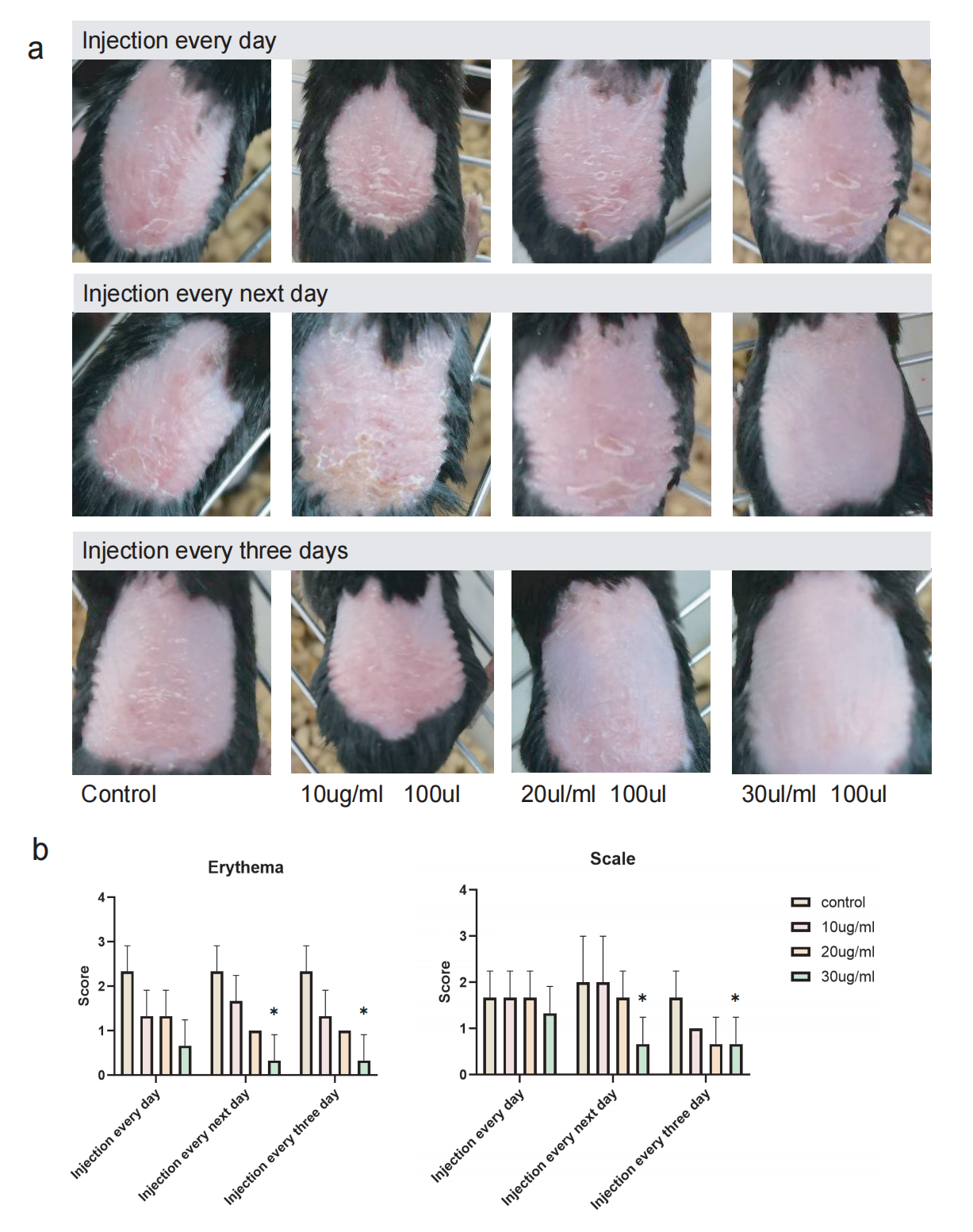

**Extended Data Fig7. Exploration of the Optimal Concentration of Secukinumab for Intervention in Psoriasis**

**a,** Mice with imiquimod-induced psoriasis were administered secukinumab at doses of 0, 10 µg/ml, 20 µg/ml, and 30 µg/ml every day, every two days, or every three days, with three mice per group;
**b,** Changes in erythema and scaling on the dorsal surface of the mice were observed following different intervention conditions.

Significance indicated as *P < 0.05, **P < 0.01, ***P < 0.001, with ns denoting not significant. Statistical analysis was performed using the t-test.

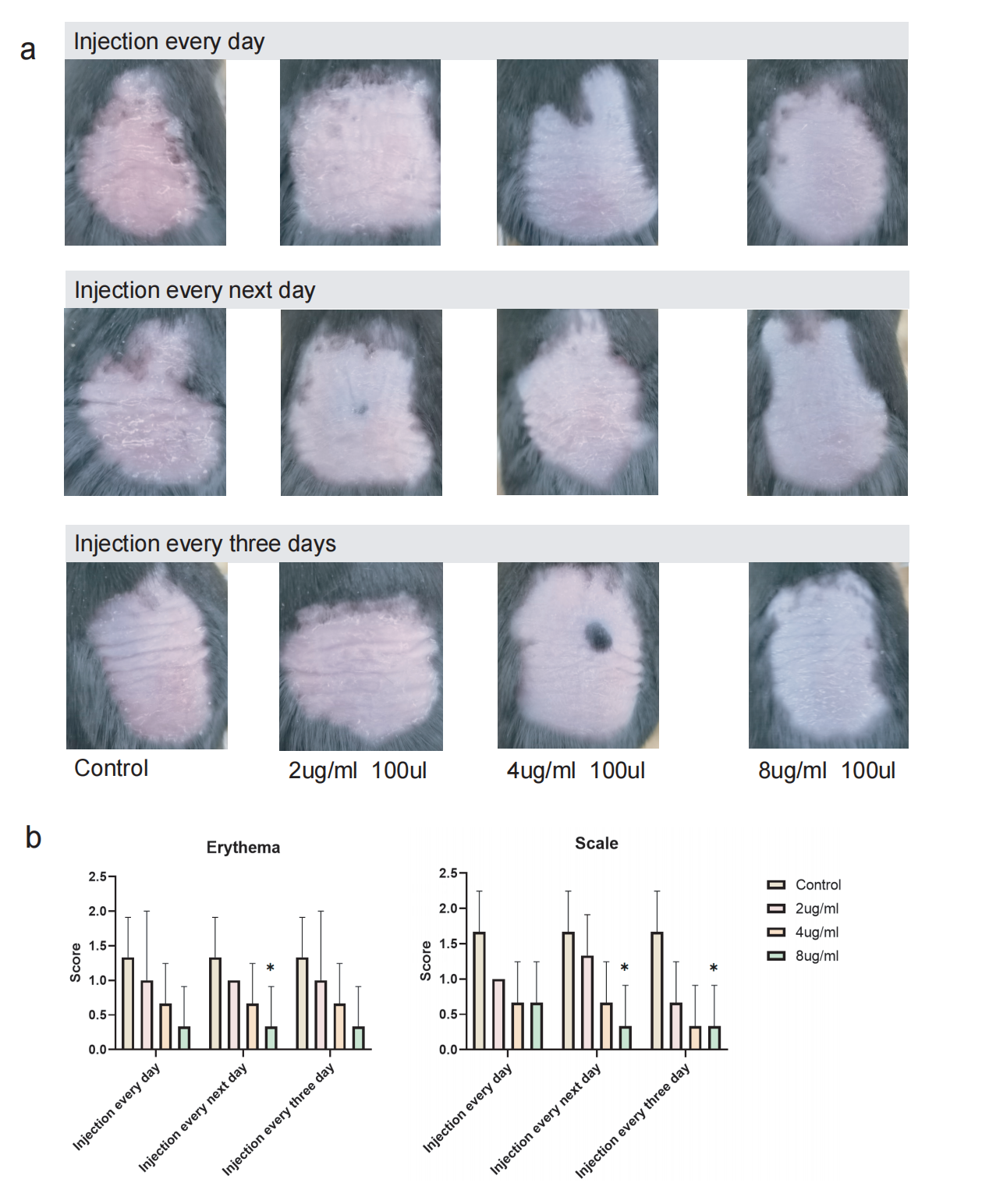

**Extended Data Fig8.Exploration of the Optimal Concentration of Ixekizumab for Intervention in Psoriasis**

**a,** Mice with imiquimod-induced psoriasis were administered Ixekizumab at doses of 0, 2 µg/ml, 4 µg/ml, and 8 µg/ml every day, every two days, or every three days, with three mice per group;
**b,** Changes in erythema and scaling on the dorsal surface of the mice were observed following different intervention conditions.

Significance indicated as *P < 0.05, **P < 0.01, ***P < 0.001, with ns denoting not significant. Statistical analysis was performed using the t-test.

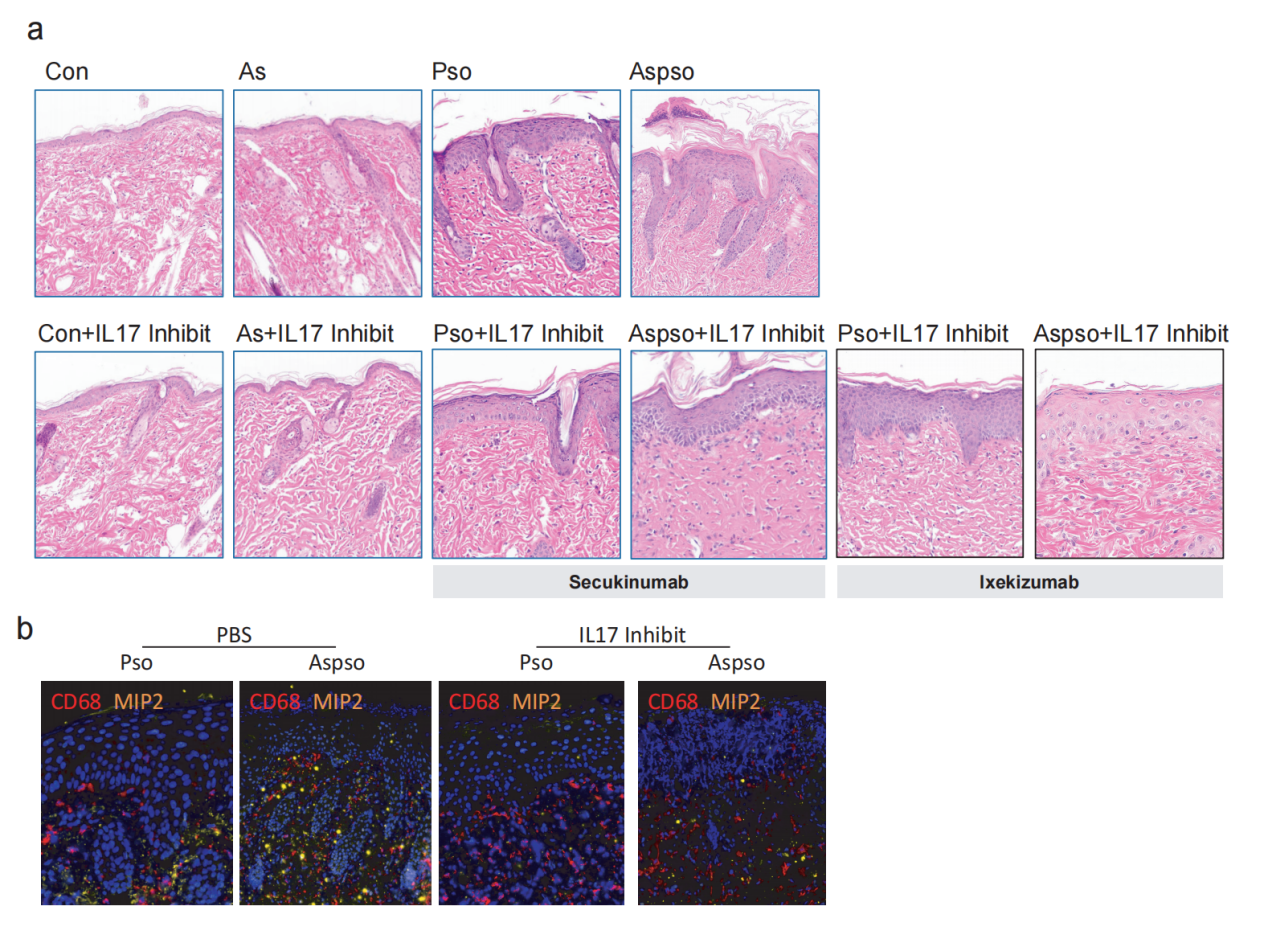

**Extended Data Fig9. Atherosclerosis Aggravates Psoriasis Progression Accompanied by Upregulation of Inflammatory Factors**

**a,** H&E staining of skin lesions in each experimental group;

**b,** Immunofluorescence showing alterations of MIP2 in each experimental group.

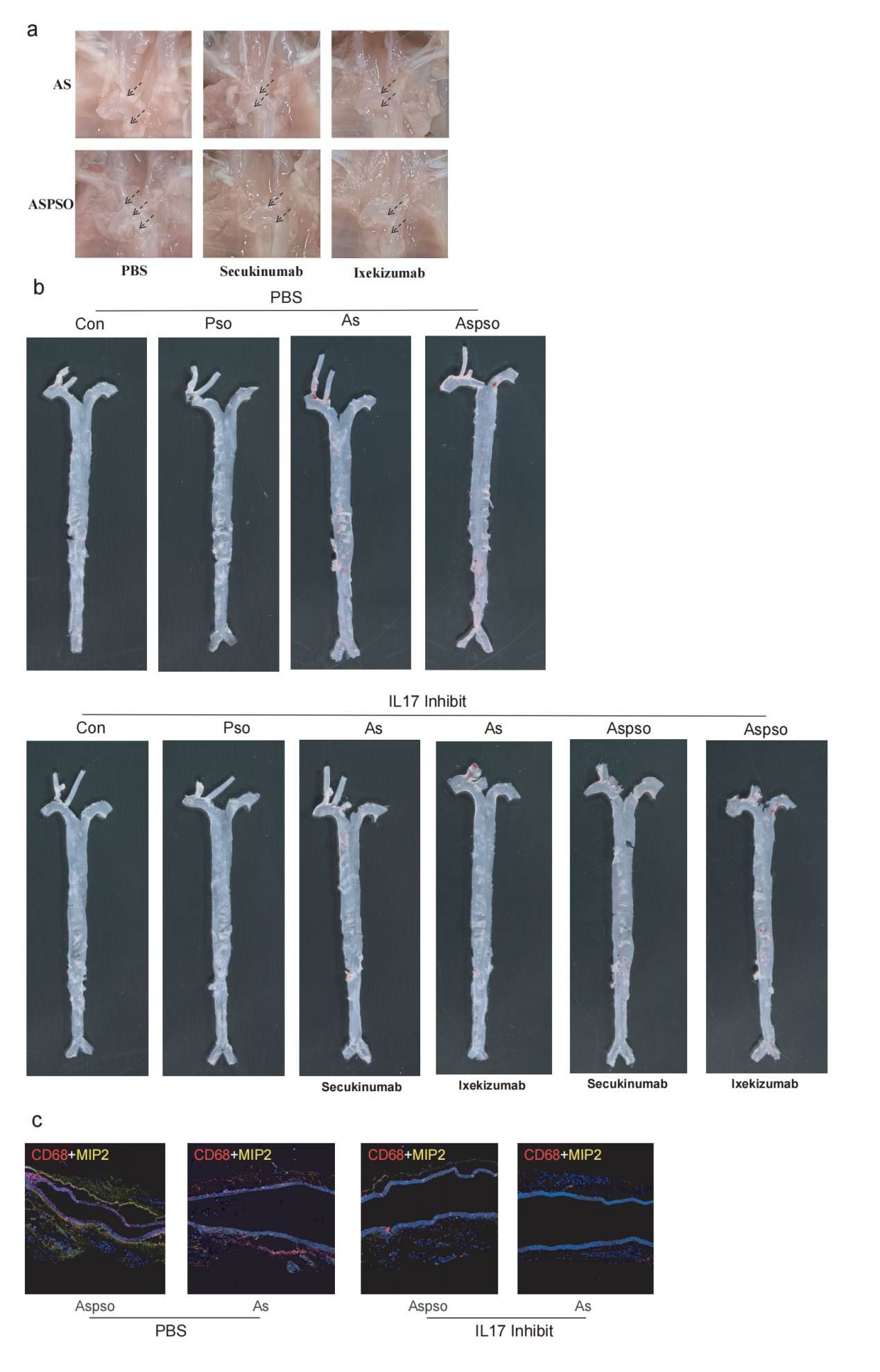

**Extended Data Fig10.** Psoriasis Exacerbates Atherosclerosis via MIP-2

**a,** Gross appearance of atherosclerotic plaque formation in the aortic arch, carotid artery, and aortic root;

**b,** Gross Oil Red O Staining of Atherosclerotic Plaque Formation in the Aorta;

**c,** Immunofluorescence Staining of the Aorta

**Extended Data Table1**

Patient characteristics of immunofluorescence

|  | Unstable Plaque | | | Stable Plaque | | |
| --- | --- | --- | --- | --- | --- | --- |
|  | P1 | P2 | P3 | P4 | P5 | P6 |
| Sex | Man | Man | Man | Man | Man | Man |
| Age,year | 78 | 69 | 72 | 60 | 71 | 67 |
| TIA attack | 0 | 1 | 0 | 1 | 0 | 0 |
| Diabetes | 1 | 0 | 0 | 1 | 0 | 1 |
| CHD | 0 | 0 | 0 | 0 | 0 | 0 |
| Hypertension | 1 | 1 | 0 | 1 | 1 | 0 |
| Smoke | 1 | 0 | 0 | 1 | 0 | 1 |

TIA:Transient Ischemic Attack; CHD:Coronary Heart Disease; 0 represents none, and 1 represents presence.

**Extended Data Table2**

[Patient characteristics](https://www.sciencedirect.com/topics/medicine-and-dentistry/patient-characteristics" \o "Learn more about Patient characteristics from ScienceDirect's AI-generated Topic Pages) for Single-cell transcriptome sequencing.

| Characteristics | Aspso1 | Aspso2 | Pso1 | Pso2 |
| --- | --- | --- | --- | --- |
| Age,year | 72 | 61 | 52 | 68 |
| Sex | Man | Man | Man | Man |
| BMI | 26.5 | 27.7 | 29.2 | 24.8 |
| Height,cm | 175 | 181 | 185 | 168 |
| Hypertension | 1 | 0 | 0 | 0 |
| Diabetes | 0 | 0 | 0 | 0 |
| Smoke | 1 | 1 | 0 | 1 |
| PASI score at baseline | 11.2 | 12.6 | 16.6 | 13.1 |
| BSA score at baseline | 31 | 29 | 29.5 | 36 |
| Duration of the psoriasis disease | 2014-2024  (10Y) | 2012-2024  (12Y) | 2009-2022  (13Y) | 2013-2023  (10Y) |

0 represents none, and 1 represents presence.

**Extended Data Table3**

Sequence information for qRT-PCR primers

| β-actin | AGCGAGCATCCCCCAAAGTT |
| --- | --- |
|  | GGGCACGAAGGCTCATCATT |
| CXCL2 | CCCAAACCGAAGTCATAGCCA |
|  | TCTCTGCTCTAACACAGAGGG |
| CXCL3 | CCCAAACCGAAGTCATAGCCA |
|  | ACCCTGCAGGAAGTGTCAAT |
| CXCL8 | GAAGTTTTTGAAGAGGGCTGAGA |
|  | ACCAAGGCACAGTGGAACAA |
| CD86 | CCTTCCTGCTCTCTGCTAACTT |
|  | TGGCCCATAAGTGTGCTCTG |
| CD163 | AAAAAGCCACAACAGGTCGC |
|  | GGTATCTTAAAGGCTCACTGGGT |
| PPARG | GCCTTAACCTCTGCTGGTGA |
|  | TCTCCGGAAGAAACCCTTGC |
| TREM2 | ACAGCATCTCCAGGGCTGA |
|  | TGCCAGAGCAGAACAAGGAG |
| CD80 | GGGAAATGTCGCCTCTCTGA |
|  | AGGTGTAGGGAAGTCAGCTTTG |
| CCL2 | CTCGCTCAGCCAGATGCAAT |
|  | TTGGGTTTGCTTGTCCAGGT |
| CCL3 | AGCCCGGTGTCATCTTCCTA |
|  | GTTGTCACCAGACGCGGT |
| APOE | TTCCCCAGGAGCCGACTG |
|  | GGGGTCAGTTGTTCCTCCAG |
| PLTP | CTTCGGGGCCCTCTTCCT |
|  | TAATGGGATCAGAGCCAGCG |
| FABP3 | CTTCCCCCTACCCTCAGGTG |
|  | CGTGGGTGAGTGTCAGGATG |
| FABP5 | ACATGAAGGAGCTAGGAGTGG |
|  | TGATGCTGAACCAATGCACC |
| ACKR1 | AACTGTCTGCACAGGGAGACT |
|  | CTCCGCCTGGAGGACATACC |
| ICAM1 | TTCCTCACCGTGTACTGGAC |
|  | CGAGAAGGAGTCGTTGCCAT |
